# Combinatorial responsiveness of single chemosensory neurons to external stimulation of mouse explants revealed by DynamicNeuronTracker

**DOI:** 10.1101/2024.09.24.614764

**Authors:** Jungsik Noh, Wen Mai Wong, Gaudenz Danuser, Julian P. Meeks

**Author notes:** These authors contributed equally to this work (Gaudenz Danuser, Julian P. Meeks).

## Abstract

Calcium fluorescence imaging enables us to investigate how individual neurons of live animals encode sensory input or drive specific behaviors. Extracting and interpreting large-scale neuronal activity from imaging data are crucial steps in harnessing this information. A significant challenge arises from uncorrectable tissue deformation, which disrupts the effectiveness of existing neuron segmentation methods. Here, we propose an open-source software, DynamicNeuronTracker (DyNT), which generates dynamic neuron masks for deforming and/or incompletely registered 3D calcium imaging data using patch-matching iterations. We demonstrate that DyNT accurately tracks densely populated neurons, whereas a widely used static segmentation method often produces erroneous masks. DyNT also includes automated statistical analyses for interpreting neuronal responses to multiple sequential stimuli. We applied DyNT to analyze the responses of pheromone-sensing neurons in mice to controlled stimulation. We found that four bile acids and four sulfated steroids activated 15 subpopulations of sensory neurons with distinct combinatorial response profiles, revealing a strong bias toward detecting sulfated estrogen and pregnanolone.

**Teaser:** DynamicNeuronTracker extracts neuronal activity from deforming 3D calcium imaging datasets using patch-matching iterations.

## Introduction

Calcium fluorescence imaging at the single-cell resolution has opened unprecedented opportunities to investigate how information is represented and processed in neural circuits of live animals [1–3]. The imaging technology enables the recording of transient rises of intracellular calcium ions for a large population of neurons at the relevant temporal and spatial resolution. The activities of thousands of neurons can now be simultaneously monitored in the brain of freely moving animals with head-mounted microscopes [4]. When combined with transgenic mouse models of Alzheimer’s disease, autism spectrum disorders, and schizophrenia, calcium imaging can support the study of brain dysfunction in diseased states at the level of neuronal activities [5–7].

To interpret the sequences of neuronal activities in calcium imaging data, it is critical to segment single neurons and extract their calcium activities. Automatic neuron segmentation is challenging because of background signals associated with blood vessels, motion artifacts, tissue deformation, etc. [8]. Various approaches have been taken for the task of neuron segmentation. Unsupervised machine learning approaches include independent component analysis [9], a generalized Laplacian of Gaussian filtering [10], and clustering with adjacent pixel correlations [11]. Model-based approaches have been implemented in Suite2P [12], CaImAn [13–15], and FIOLA [16]. Recently, supervised deep neural network (DNN) models were developed for two-photon calcium imaging [17–20] and widefield imaging [21]. The DNN models’ applicability would depend on the model’s ability to perform well for unseen imaging datasets that can be quite different from the training datasets used to build the models. One of the widely used neuron segmentation methods is CaImAn, which utilizes constrained nonnegative matrix factorization to extract spatial neuron masks and temporal signal fluctuations simultaneously. CaImAn 3D is currently the only method that processes bona fide 3D calcium imaging data.

All discussed segmentation methods except one DNN model [20] assume and generate static neuron masks, that is, fixed regions-of-interest (ROIs). Since the assumption is often violated, image registration or motion correction for raw imaging data is a necessary preprocessing step [8]. However, calcium image registration can be almost impossible in some imaging modalities due to the lack of consistent background signals and the nature of calcium signals being intermittently active. In an extreme scenario where the whole nervous system of freely moving *C. elegans* was imaged in 3D, sparsely active neurons could not be registered, so another fluorescence marker of the cell body was imaged for segmentation purposes [2, 22]. In our previous study using 3D light-sheet imaging of the mouse vomeronasal organ (VNO) [23], we observed linear and non-linear tissue deformation during an hour of imaging that persisted after application of existing volumetric registration tools. These uncorrectable artifacts dramatically slowed efforts to manually segment active neurons, and prevented the application of existing software tools based on fixed ROI segmentation methods.

Here we introduce a neuron segmentation method named DynamicNeuronTracker (DyNT) that generates dynamic ROIs for single neurons in 3D calcium imaging of deforming tissues. Our algorithm employs exhaustive iterative optimization with local images of active neurons, achieving high accuracy in tracking neurons with highly dynamic fluorescence intensities and unstable positions. We demonstrate that the dynamic ROIs generated by DyNT accurately track densely populated neurons, while fixed ROIs generated by CaImAn are often erroneous under positional jitter. We illustrate the differences in performance between DyNT and CaImAn compared to manual segmentation. We complement the DyNT pipeline with statistical analysis of the association between the neuronal calcium activities and stimulation events for the case when the imaging is performed while controlled stimulation is applied to the neurons. To do this, the DyNT pipeline includes a series of statistical testing procedures to determine the combinatorial responsiveness of each segmented neuron to an applied sequence of randomized, interleaved blocks of stimulation, generating comprehensive lists of neuronal response profiles to stimulation input.

DyNT can be used to segment and evaluate neural activity patterns to any list of stimuli, but was used here in the context of understanding how external chemical information is converted to neural activity patterns by vomeronasal sensory neurons (VSNs) in the mouse VNO. The VNO in mammals is a sensory tissue specialized for the detection of environmental nonvolatile chemicals, including many pheromones [24]. The VNO is a mosaic tissue, populated by tens of thousands of VSNs, each of which typically expresses just one vomeronasal receptor (VR) out of a family of ∼300 unique genes [25, 26]. VRs have highly overlapping chemical sensitivities, and it is typical for VNO ligands to activate multiple VRs, and each VR is capable of detecting multiple ligands [27]. VSNs, like other olfactory populations, thus encode chemical information using a combinatorial strategy [28–31]. The combinatorial encoding strategy supports the identification of patterns of ligands in natural blends, providing a mechanism by which chemical origins (*e.g.* animals of a particular species, sex, maturity, health, etc.) are computed at the population level.

Only a small fraction of ligand-receptor mappings has been discovered so far in VSNs, and even less is known about the full chemosensory repertoire encoded by the VNO. Previous studies have explored chemosensory mechanisms for monomolecular steroid ligands, including sulfated steroids present in estrus mouse urine [31–33], and bile acids present in mouse feces (cholic, deoxycholic, lithocholic, chenodeoxycholic acids; CA, DCA, LCA, CDCA, respectively) [23, 34]. In this study, we analyzed 3D calcium imaging datasets of VNO stimulated by four bile acids and four sulfated steroids. We examined 1,628 neurons segmented by DyNT, and determined their combinatorial responses. We identify 15 subpopulations among the segmented VSNs that show distinct combinatorial response profiles to the applied 8 ligands. These results identify disproportionate representation of chemosensory information related to a sulfated estrogen (E0893) and pregnanolone (P3865), suggesting current models of vomeronasal chemosensation may be incomplete. Overall, these data demonstrate the power of DyNT for extracting neuronal activity patterns from dynamic neurons with unstable positions, and for quantifying the encoding of chemical information by a population of stimulated chemosensory neurons.

## Results

### Uncorrected spatial displacement disrupts neuron segmentation via fixed ROI methods

In a previous study, we imaged functionally intact VNOs to investigate ligand-receptor interactions for pheromone-sensing neurons [23]. VNOs were dissected from mice expressing genetically encoded calcium indicator GCaMP6s, and a volume of the epithelium with an approximate size of 0.7 × 0.7 × 0.4 mm^3^ was imaged into one field-of-view (FOV), permitting a temporal sampling of 3 sec/frame (0.33 Hz) for 75 min. During imaging, the tissue was stimulated with 8 known monomolecular ligands: four bile acids (CA, DCA, LCA, CDCA) and four sulfated steroids consisting of an androgen (A6940), an estrogen (E0893), a pregnanolone (P3865), and a glucocorticoid (Q1570). The panel of stimuli also accounted for varying concentrations (each bile acid at 1 µM and 10 µM), natural blends of social odorants as positive controls (dilute female/male mouse feces, FF/MF), and the vehicle (negative) control (Ringer’s), for a total of 15 stimuli that were presented to tissues 5 times each in a randomized, interleaved block design. Upon stimulation, VSNs displayed seemingly selective responses as previously observed [23].

The volumetric movies displayed slow non-linear deformation of the VNO tissue during an hour of imaging, as well as fast uncorrected translational artifacts (Videos S1 and S2). Because of the slow persistent tissue deformation, the locations of single neurons drifted throughout the recordings (Figure S1A). Manual extraction of activation time series by tracking individual neurons in these image time-lapse sequences was extremely laborious, and introduced the possibility of human bias during ROI selection [23].

To establish a robust, automated pipeline for the extraction of activation time series, we first tested if other existing registration methods could compensate for the observed uncorrected tissue deformation. We applied a rigid registration based on phase correlations and a non-rigid registration tailored to motion correction for calcium imaging data [35]. Substantial positional jitter remained after either of the registrations (Videos S3 and S4). In particular, we observed numerous instances where firing neurons were significantly displaced between consecutive time points (Figure 1A, Figures S1B-S1D) preventing the extraction of static neuron masks. Because of the lack of background signals, further refinement of the registration by a more granular, yet still continuous, deformation field seemed unpromising. Instead, we decided to develop a segmentation of individual neurons by computing dynamic ROIs.

**Fig. 1.**
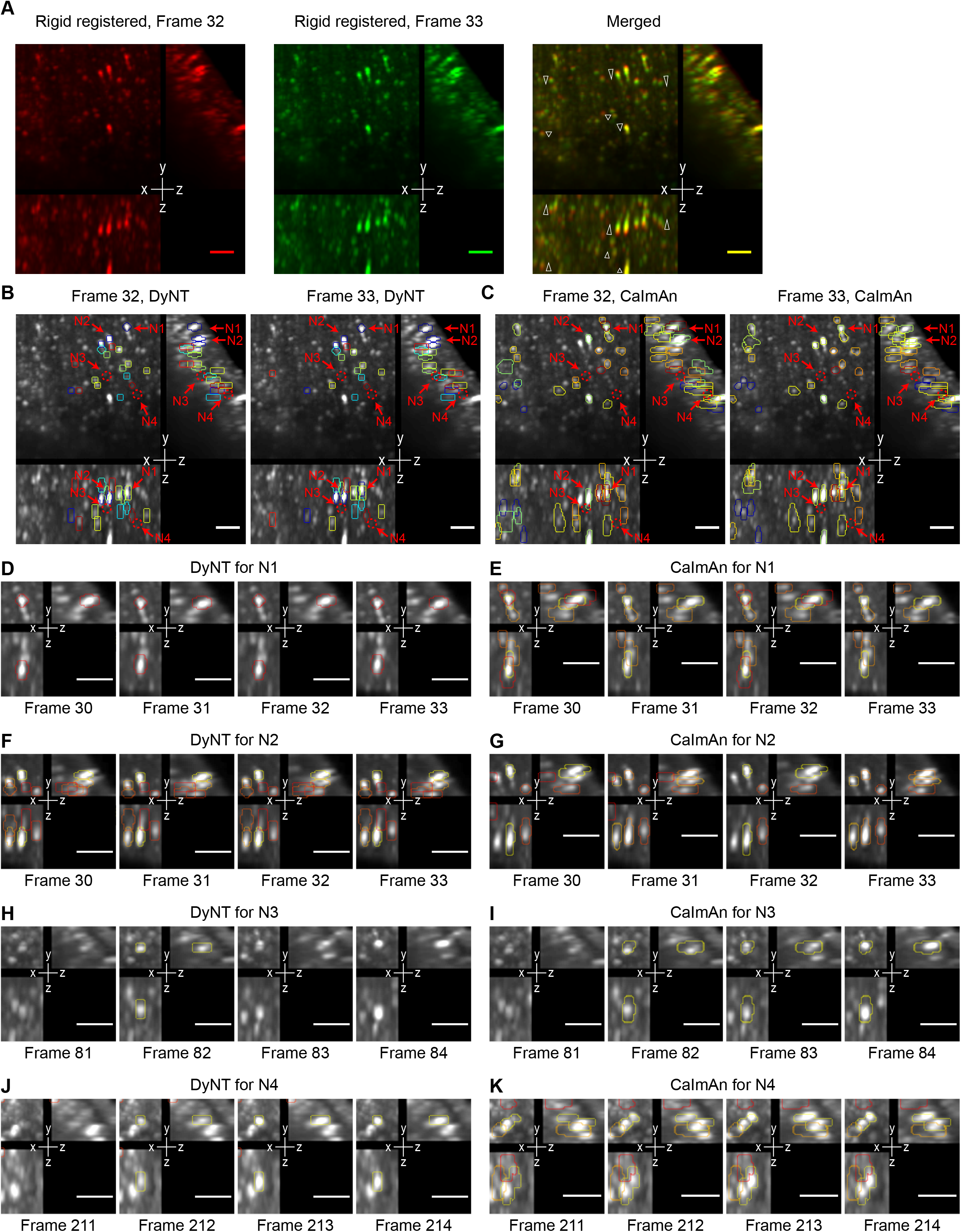
Jittering neurons in deforming tissues and their segmentation using dynamic and fixed ROI methods. (**A**) Positional jitters between two consecutive time points are highlighted for several neurons (white triangles) on maximum intensity projections (MIPs) of sub-volume images after rigid registration. All scale bars, 50 µm. (**B** and **C**) Active neurons segmented by DyNT (**B**) and CaImAn (**C**) (dynamic and fixed ROI methods, respectively). Locations of example neurons magnified in (**D**-**K**) are indicated. (**D** and **E**) Local MIP images around neuron N1. ROI extracted by DyNT is red in (**D**); CaImAn generates two ROIs (red and yellow) (**E**). The neuron moves down in y-axis by 1-2 pixels at frame 31 and moves back up at frame 32. ROIs with different colors indicate masks for neighboring neurons throughout (**D**-**K**). (**F** and **G**) Local MIP images around neuron N2. ROI extracted by DyNT is yellow in (**F**); CaImAn generates two ROIs (yellow and orange) (**G**). (**H** and **I**) Local MIP images around neuron N3. N3 is the brightest at frame 82 and an adjacent neuron becomes the brightest at frame 84. ROI extracted by DyNT is yellow in (**H**). The ROI by CaImAn (yellow in (**I**)) includes the adjacent active neuron. (**J** and **K**) Local MIP images around neuron N4. N4 is active in frames 212-214 and segmented by DyNT accordingly; yellow ROI in (**J**). ROI by CaImAn shown as yellow in (**K**) is active in frames 211-214 because it includes adjacent active neurons.

Figure 1 highlights the differences between fixed and dynamic ROI methods in the presence of positional jitter. It provides a head-to-head comparison of DyNT and CaImAn 3D in a sub-volume (∼7%) of the full FOV of one of the VNO calcium time-lapse sequences. ROIs are shown only when they contain a significant calcium activity. Unlike DyNT, CaImAn segmentation includes abnormally large ROIs (Figures 1B and 1C). Many of the large ROIs appear across all time points (Video S5). The number of these large ROIs increased when we attempted to segment more neurons by adjusting a corresponding parameter in CaImAn. This suggests that positional jitter confuses the fixed ROI segmentation, especially for spatially close neurons. For example, we found numerous instances where a single neuron was captured by two different fixed ROIs, while a corresponding dynamic ROI of DyNT correctly followed the small movements (Figures 1D-1G, Videos S6 and S7). In other instances, we found that spatially adjacent jittering neurons were segmented into one fixed ROI of CaImAn, while DyNT separated them accurately (Figures 1H-1K, Videos S8 and S9). This is a critical feature of DyNT, because the attribution of two individual neurons’ calcium activity to a single neuron introduces false stimulus tuning properties that may cause improper conclusions to be drawn from the data. These instances demonstrated that fixed ROI segmentation is error-prone for dynamic, deforming, and/or incompletely registered calcium imaging data.

### Tracking dynamically active neurons in dynamic, deforming 3D calcium imaging data

To track and segment the dynamically active, spatially jittering neurons, DyNT first detects active neurons as small, bright objects and tracks their movements within consecutive time frames. Next, DyNT groups the firing events throughout the time frame that belong to the same neuron (Figure 2A). Effectively, this first task is to perform 3D particle tracking for all detected point sources (transient increases in GCaMP6s fluorescence) in the 3D movie. DyNT utilizes u-track 3D [36] to identify tens of thousands of neuronal firing events. To track neurons with different sizes, DyNT employs multi-scale particle detection (see Methods).

**Fig. 2.**
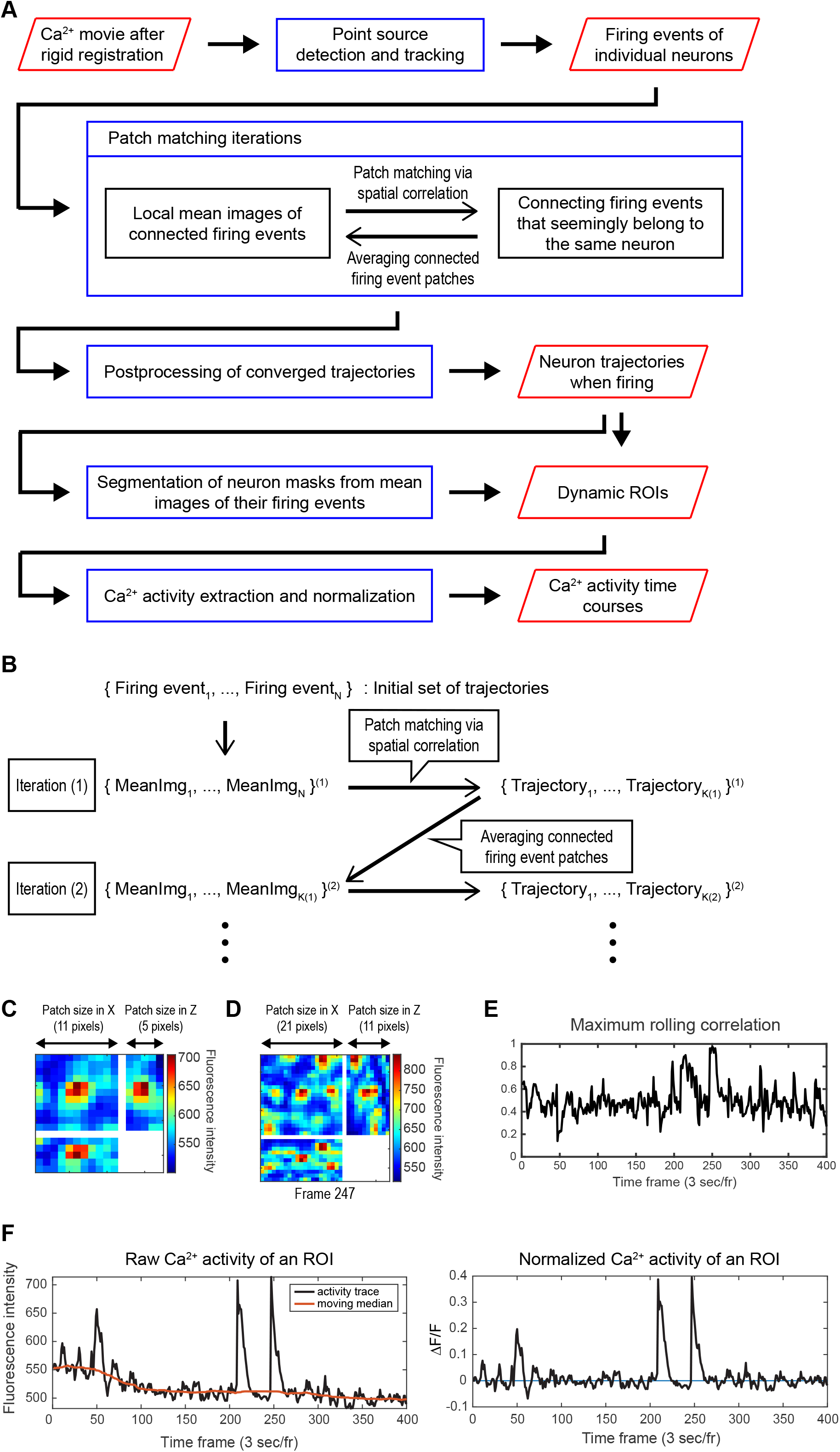
Overview of DynamicNeuronTracker. (**A**) Workflow of DynamicNeuronTracker (DyNT) that takes a rigid-registered calcium video as input and produces dynamic ROIs and their calcium activity time courses. Red boxes indicate data objects and blue boxes indicate computational operations. (**B**) Patch matching iterations in (**A**) are illustrated. In the first iteration, a local mean image of a neuron firing event is matched to another firing event nearby at different time frames using spatial correlations as illustrated in (**C**-**E**). From a resulting trajectory, we obtain an updated local mean image utilized to run the second iteration. (**C**) An example MIP of a local mean image of a firing neuron. The 3D patch size (11 x 11 x 5 pixels) is a user input. (**D**) An example MIP of a neighborhood local volume that contains a neuron being tracked. Rolling spatial correlations between a local mean image (**C**) and its neighborhood volume (**D**) are computed in 3D to examine when the mean image appears again at late time points. The size of the neighborhood volume (21 x 21 x 11 pixels) is also a user input depending on the degree of jittering. (**E**) Maxima of rolling spatial correlations at every time frame. High correlations indicate the neuron is likely to be active at the time. (**F**) (Left) Raw calcium activities of an ROI are obtained by averaging fluorescence intensities within the ROI at every time frame (black). Red indicates moving medians of the raw activities computed in a rolling window of 81-frames. (Right) Normalized calcium activities, ΔF/F = (raw activity – moving median)/moving median.

For the second task, DyNT iteratively connects the detected firing events for individual neurons using correlation-based patch-matching (Figure 2B). The patch-matching iteration starts by computing the local mean image of each firing event (Figure 2C). For a given firing event, we consider a wider local volume containing the event (Figure 2D) and examine when the mean image appears again across all the time frames using rolling spatial correlations (Figure 2E). If the maximum of the rolling correlations at any time point is higher than a user-specified threshold, then the corresponding neuron is set as firing at that time point (see Methods). The locations of rolling correlation maxima define the displacements of the firing neurons. These lead to preliminary information on when and where each neuron is firing, which allows us to update the local mean images of the neurons when firing. Thus, spatially jittering neuron trajectories and their mean images of firing events iteratively update one another. After several iterations, the individually detected firing events of a single neuron are thus connected into a single neuron trajectory (Figure 2B).

After the patch-matching iteration, a 3D Gaussian density function is fitted to each local mean image of the neurons, and its central mass defines a mask for the neuron. Each mask combined with the neuron trajectory results in a dynamic ROI. Because the neuron trajectory is only defined when the neuron is firing, the ROI location when the neuron is inactive is set to the location of the last activation. Averaged fluorescence intensities within the dynamic ROIs across the time frames are collected as raw calcium activities of identified single neurons, which are normalized by using rolling medians as the baseline fluorescence (Figure 2F).

### Improved measurement of single-neuron calcium activities using spatial correlations

Visual inspection indicated that DyNT segmented approximately 25% - 50% of all the visually detectable neurons (Figure 3A, Video S10). To validate the accuracy of the segmentations, the DyNT pipeline created maximum intensity projection (MIP) videos of local volumes around every segmented dynamic ROI. To focus the visual inspection on time points of neural firings, the pipeline classified the calcium activities of each neuron into high and low states using K-means clustering and produced the local volume videos displaying only when the ROI is active (see Videos S6-S9).

**Fig. 3.**
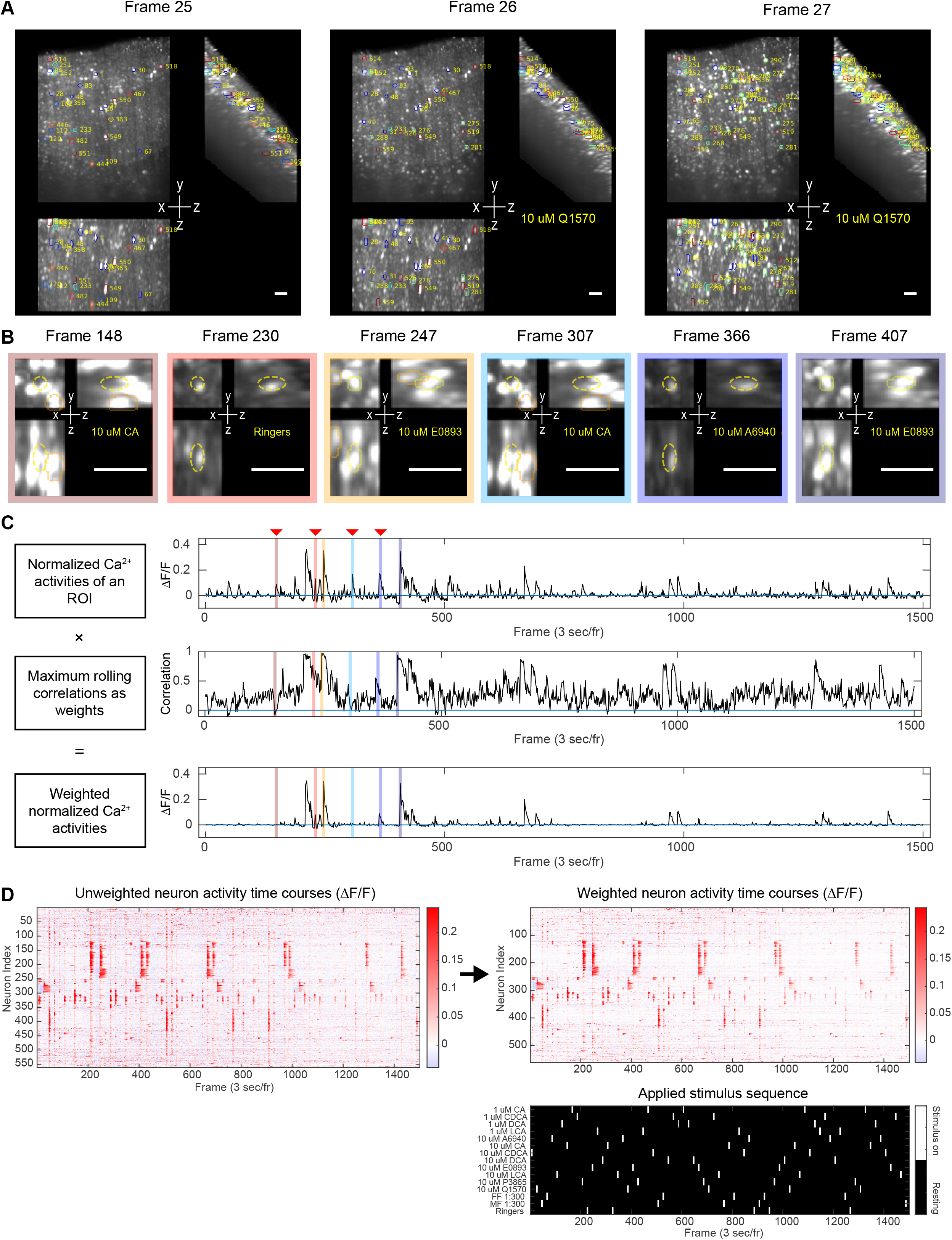
Extraction of neuronal calcium activities using a weighting scheme with spatial correlations. (**A**) Segmented dynamic ROIs overlaid to consecutive MIP images of an entire VNO tissue when they are active. Different ROIs are numbered and color-coded. The stimulation status is reported at the lower right. All scale bars, 50 µm. (**B**) MIP images of a local volume around a target neuron at different time points. The target neuron is active (yellow contour) at time points 247 and 407, and inactive (dashed yellow) at other time points. The activities of adjacent neurons infiltrate the target ROI when the target neuron is inactive. (**C**) A weighting scheme to improve calcium activity measurement. (Top) Normalized calcium activities of the target neuron in (**B**) are shown to have spurious peaks, caused by image interference from adjacent neuronal activities. Frame numbers are indicated by red triangles. The six time points in (**B**) are marked with the matched color. (Middle) The maximum rolling correlation time course is the one obtained after patch matching iterations for the target neuron are finished. (Bottom) The multiplicative weighting reduces many spurious peaks and noises across the time frames. (**D**) (Top) Unweighted and weighted normalized calcium activities of 561 neurons segmented from the VNO video shown in (**A**). (Bottom) Schedule of randomly repeated 15 different stimuli during the imaging period.

While visually validating the dynamic ROIs and their calcium activity time courses, we observed that adjacent neuronal activities interfered with the signal of the target ROI, producing spurious peaks in the calcium traces (Figures 3B and 3C). The interference occurred also at time points in which the target neuron was not active, altering the statistical testing of neuronal responsiveness to given stimuli. The interference could not be eliminated by simply making smaller ROIs.

To eliminate spurious activity peaks, we introduced a weighting scheme when collecting neuronal activities via the ROIs. To track the jittering neurons, DyNT had already calculated the maximum rolling spatial correlations, which provided information about the likelihood of the target neuron being active at each time point. The maximum spatial correlations for each neuron, after negative correlations were truncated, were used as multiplicative weights for the normalized calcium activities (Figure 3C, See Methods). The weighting scheme significantly reduced the spurious activity peaks and noise (Figure 3C). We visualized the unweighted and weighted normalized calcium activities of an entire video together with a known sequence of randomized, interleaved stimulus trials (Figure 3D). This apposition of stimuli and response schedules provides a broad view of the qualitative correspondence between individual stimuli (sulfated steroids, bile acids, and natural blends of chemosignals) and neural activation.

### Performance evaluation with reference to manual segmentation

To quantify the neuron segmentation performance of DyNT, we compared DyNT and CaImAn in relation to manually segmented ROIs. Using DyNT and CaImAn, we processed three VNO calcium videos, for which manual segmentation has been already completed [23]. The manual segmentation generated fixed volumetric ROIs for well-registered VSNs that reliably responded to given stimuli. It could not segment densely populated regions, or regions with uncorrected spatial displacement. It was additionally biased to segment stimulus-responsive neurons, whereas DyNT and CaImAn segmented both stimulus-specific and spontaneously active neurons without priors (Figures 4A and 4B). Thus, the manually segmented fixed ROIs do not represent an absolute ground truth for performance evaluation. Nonetheless, we computed recall performances for DyNT and CaImAn given the manually segmented masks.

**Fig. 4.**
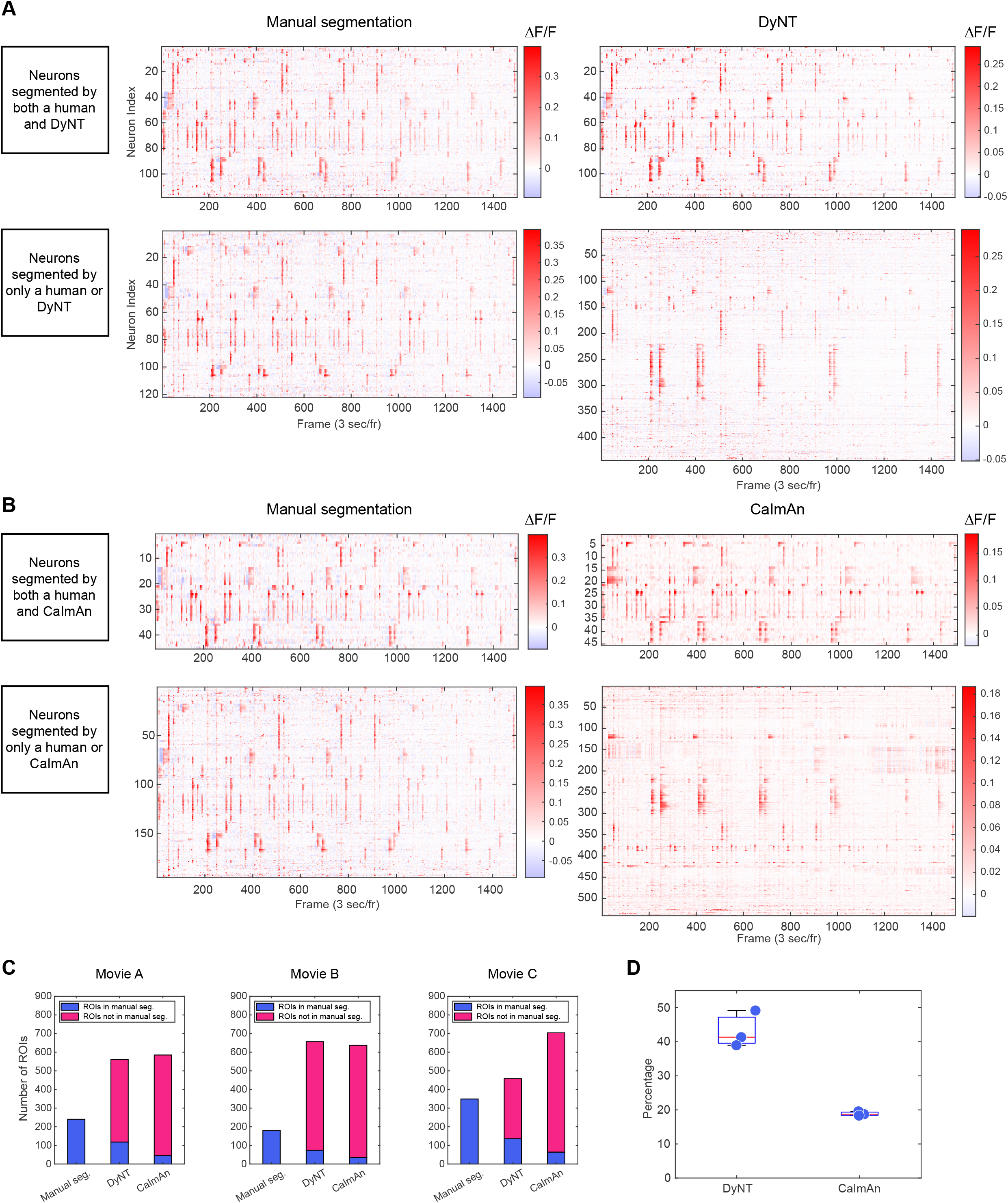
Performances of DyNT and CaImAn in reference to manual segmentation. (**A** and **B**) Comparison of segmented ROIs’ calcium activities (ΔF/F) between manual annotation and DyNT (**A**) (and CaImAn (**B**)). If the Jaccard index between two ROIs is greater than 0.3, the two ROIs are determined to segment the same neuron. (Top) Activities of the neurons segmented by both manual annotation and DyNT (**A**) (CaImAn (**B**)). (Bottom) Activities of the neurons segmented by only manual or DyNT. (**C**) The number of neurons segmented by manual annotation, DyNT and CaImAn is visualized across three VNO videos. The number of neurons recaptured by DyNT and CaImAn among manually segmented ones is presented in blue. Magenta represents the neurons segmented by only the algorithms not by the manual annotation. (**D**) The recapturing ratios for three VNO videos.

To define the association between manually- and machine-segmented masks, we utilized a Jaccard similarity index > 0.3 as a threshold. For dynamic ROIs, we averaged the Jaccard indexes over all time points. Figures 4A and 4B directly illustrate the correspondences between manual segmentation and DyNT vs CaImAn (manual: 240 ROIs, DyNT: 561 ROIs, CaImAn: 585 ROIs). DyNT ROIs included 118 (49%) out of the 240 manual ROIs, while CaImAn ROIs included 45 ROIs (19%). Over three VNO videos, DyNT showed a much higher recall compared to CaImAn (Figures 4C and 4D).

### Processing large-volume videos using parallel segmentation

Single-neuron segmentation for calcium imaging data is a task that can be easily divided into multiple parallel operations. The computation of a single-neuron ROI is independent of the pixel intensities distant from the ROI. The DyNT pipeline supports parallel segmentation by dividing an original volume into 8 (or more) sub-volumes of equal size. Because dynamic ROIs near the boundary of a sub-volume may be partially contained, the sub-volumes are generated allowing overlaps. The sub-volume videos can be processed in parallel using multiple computing nodes, if available. This parallelization also allows users to overcome computer memory limits when processing large-volume videos.

After each sub-volume is processed, the DyNT pipeline integrates sub-volume outputs into the full-volume output, while assessing potentially redundant ROIs within overlapping areas and merging the redundant ROIs into unique ones without requiring an additional parameter. This parallelization is scalable since a sub-volume video can be further parallelized in the same way, that is, 64 sub-volume videos for an original video. Thus, the DyNT pipeline provides the capacity to process large-volume videos via its own parallelization functionality.

### Comprehensive segmentation output reveals differential chemosensory functions of vomeronasal sensory neurons

Each VSN generally expresses one dominant VR. Given that each VR can detect one or multiple ligands, the calcium activity pattern of an individual VSN to a panel of stimuli represents a signature of the ligands sensed by the expressed receptor [28]. To determine which applied stimuli activated individual VSNs, we first analyzed the calcium activity of 561 VSNs segmented from the 3D movie shown in Figure 3D. Using one-sided t-tests and false discovery rate (FDR) control [37] (See Methods), we statistically tested whether each neuron’s responses to each stimulus were significantly higher than its responses to Ringer’s, which is referred to as the marginal responsiveness hereafter. Then, the responsiveness across different stimuli was combined to determine the subset of stimuli an individual neuron was activated by, which is referred to as a combinatorial responsiveness profile. For example, Figures 5A and 5B (Video S11) show the MIP images and short-term calcium activity curves of a VSN (ROI 303) that responded to only CA. Of note, this VSN consistently fired only for 10 µM CA, not for 1 µM CA or for 10 µM DCA. Figures 5C and 5D show the MIP images and activity curves of a VSN (ROI 344) that responded to only DCA (Video S12); and Figures 5E and 5F those of a VSN (ROI 156) that responded to both E0893 and P3865 (Video S13). In this case, there was a DCA-responsive VSN adjacent to the target neuron, as seen in frame 288 (Figure 5E, Video S13). The activity of this adjacent DCA-responsive neuron interfered with ROI 156, leading to statistically significant responses of ROI 156 to 10 µM DCA under a t-test (Figure 5F). However, the responses to DCA did not reach significance at the FDR level of 10%, resulting in a responsiveness profile that aligned with our visual inspection. While this example highlights the robustness of our pipeline in eliminating the effects of spurious activity peaks and other sources of noise, it also shows that the proposed weighting scheme retains a risk of signal leakage between closely neighboring neurons. The remaining interference represents one of several noise sources in the neuronal activities extracted from the imaging data and segmented ROIs (Figure 5F).

**Fig. 5.**
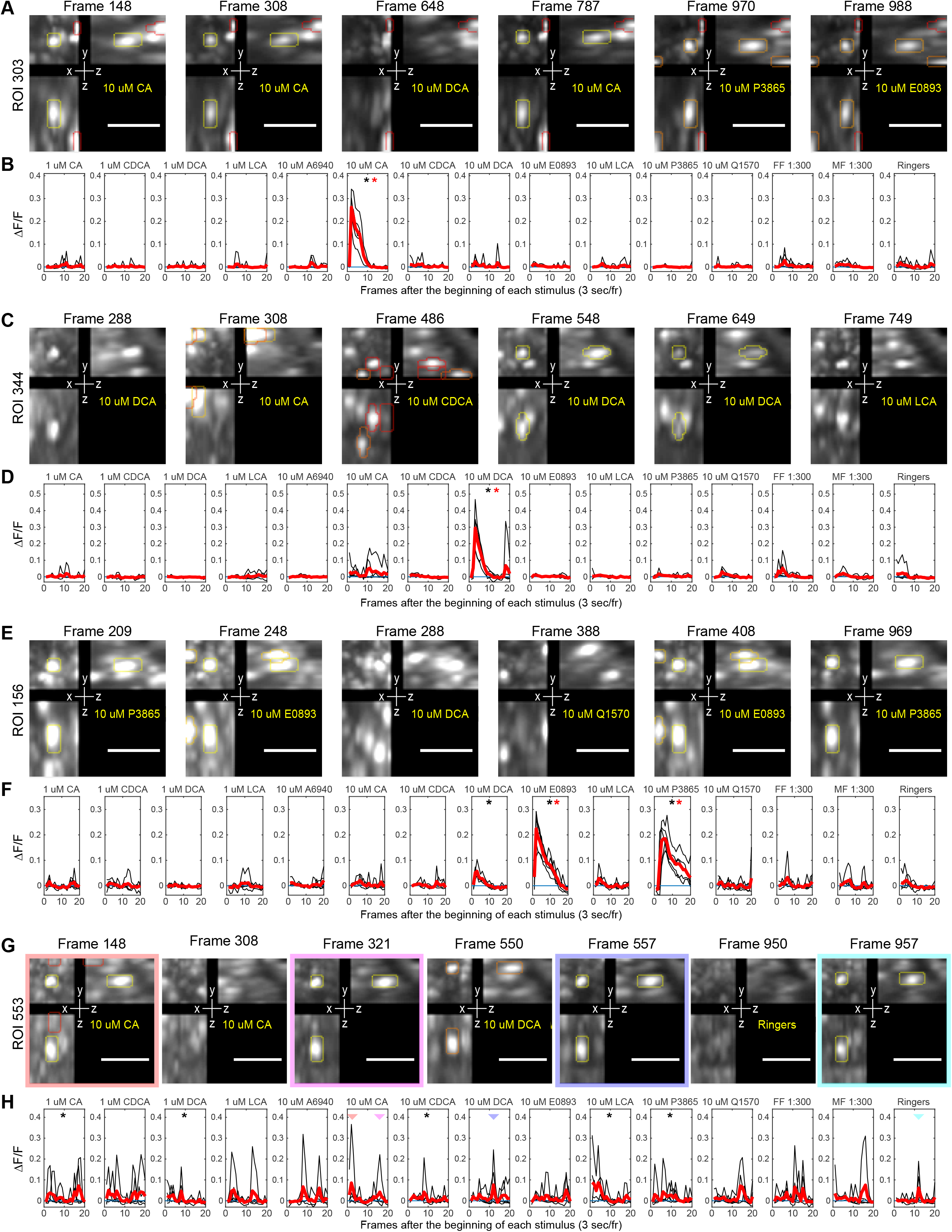
DyNT output displays selective responses of VSNs to bile acids and sulfated steroids. (**A**, **C** and **E**) Representative local MIP images of a neuron that exclusively responded to CA, DCA, and E0893 and P3865, respectively. Yellow masks indicate the target neuron being active. Masks with different colors indicate other neurons throughout **A**, **C** and **E**. The stimulation information is provided at the lower right. All scale bars, 50 µm. (**B**, **D** and **F**) Calcium activity response curves of the neurons shown in (**A**, **C** and **E**), respectively, for each stimulus (black). Each trial period is 20 frames (60 s), with each stimulus delivered during the first five frames (15 s). The red curve indicates an averaged response curve over five trials for each stimulus. The black star indicates the across-trial responses for a stimulus are significantly higher than the ones for Ringer’s (P < 0.05, one-sided t-test). The red star indicates the significance after FDR control (adjusted P < 0.1). (**G**) Local MIP images of a representative neuron that is spontaneously active. (**H**) Calcium activity response curves of the spontaneously active neuron (**G**). Active MIP images are outlined in (**G**), with corresponding time points indicated by triangles with matching colors in (**H**).

Approximately 30% of the ROIs segmented by DyNT did not show an obvious, time-locked response to any of the presented stimuli (Figure 3D). This is expected, as only a subset of the dozens of known ligands for these receptors was presented. This is a feature of DyNT, not a deficiency, as VSNs are known to be spontaneously active, and their spontaneous activation properties are an important consideration for neural coding [38]. DyNT thus overcomes user bias in selecting ROIs – a significant drawback of manual annotation of chemosensory-driven activity – and enables the discovery and analysis of neural populations with interesting features aside from their chemosensory tuning. For example, in this dataset, we identified VSNs, including the neuron segmented in ROI 553 (Figures 5G and 5H, Video S14) that showed a pattern of consistent activation many seconds after stimulus offset, including the negative control Ringer’s solution. The function of such neurons is unknown but might be involved in providing additional information relevant to this sensory system.

### Statistical procedures to determine the combinatorial responsiveness of single neurons to a given set of stimuli

We applied the same statistical activation profiling to a total of 1,628 VSNs segmented from all three of the available VNO videos to determine the combinatorial responsiveness. We compared their across-trial responses to 14 different stimuli with the responses to Ringer’s (Figure 6A). After t-testing and subsequent FDR control, we determined the marginal responsiveness of individual VSNs (Figure 6B). This statistical step is equivalent to testing and constructing the edges of a graph between a set of 14 stimuli and a set of 1,628 neurons (Figure 6C). It is worth noting that the marginal responsiveness outcomes can contain up to 10% false positives.

**Fig. 6.**
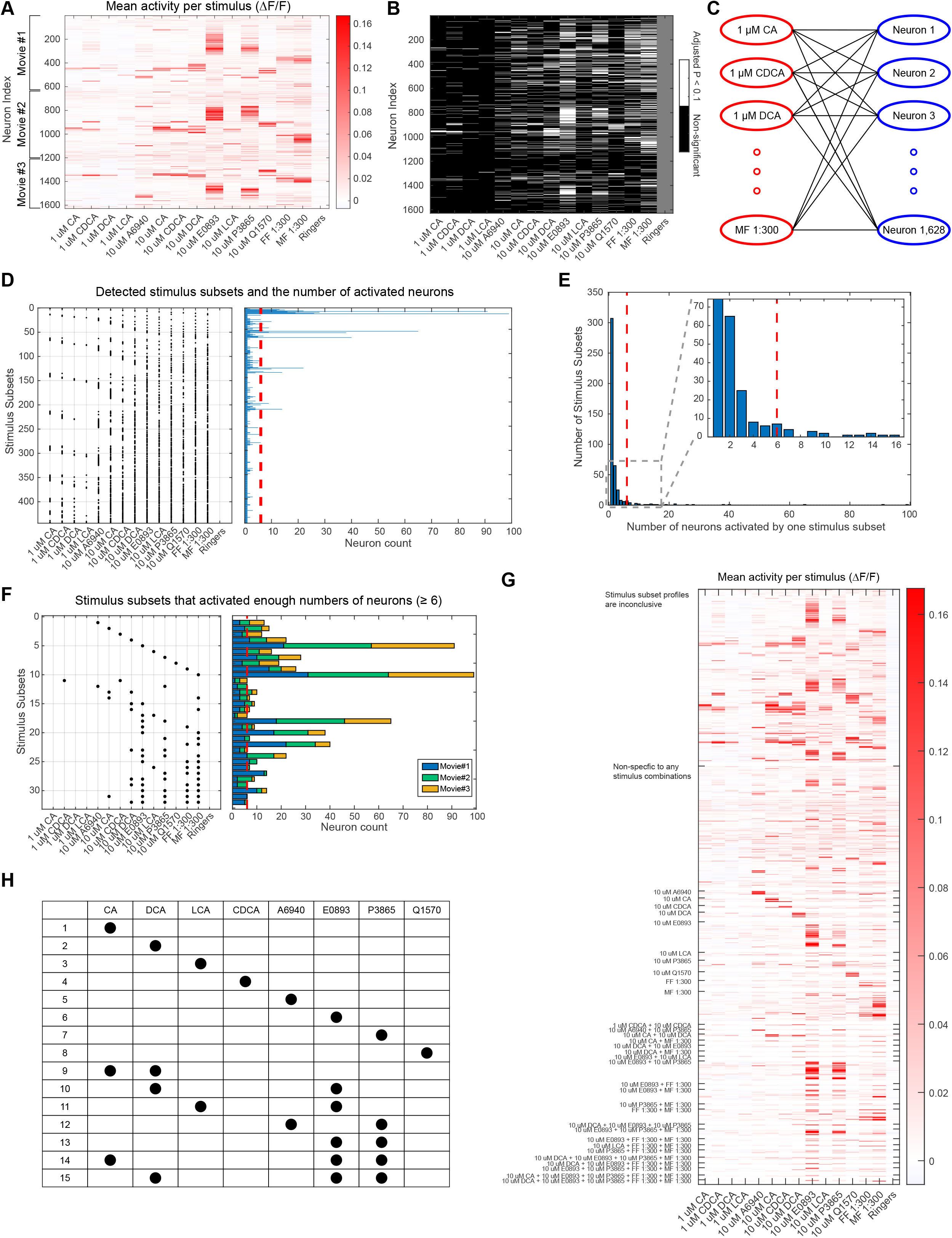
DyNT determines the combinatorial responsiveness of VSNs and identifies ligand subsets that activate distinct neuronal subpopulations. (**A**) Averages of calcium activity responses (ΔF/F) over different trials for individual stimuli and 1,628 neurons segmented from three VNO videos. (**B**) Adjusted P-values after marginal comparisons of the mean responses under a stimulus and Ringer’s condition using t-tests and FDR controls. (**C**) A representation of the statistical step in (**B**) as a graph reconstruction problem. Each significant adjusted P-value (< 0.1) leads to drawing an edge between a stimulus and a neuron in the graph, but up to 10% of the drawn edges can be false positives. (**D**) From the adjusted P-value matrix in (**B**), subsets of significant stimuli for individual neurons are collected and visualized in dot representations. The numbers of neurons which lead to those stimulus subsets are counted and visualized. The red line indicates a threshold for the neuron counts derived from (**E**). (**E**) A histogram of the number of neurons that are activated by significant stimulus subsets in (**D**). (Inset) A magnified histogram when the activated neuron counts are small. It displays an exponential decay of the number of stimulus subsets at the beginning, which is the typical distribution pattern of random events. But there is a peak at the neuron count of six, which is a deviation from the random event pattern. The red line indicates a threshold for the neuron counts determined by the peak. (**F**) 32 stimulus subsets and the number of corresponding activated neurons after excluding the stimulus subsets that activated only a small number of neurons (< 6) from (**D**). The threshold six is chosen from the histogram (**E**). Movie indexes of the activated neurons are color-coded. It shows that most of the combinatorial responsiveness profiles are reproduced over the three movies. (**G**) Per-stimulus mean responses of segmented neurons visualized after determining their combinatorial responsiveness. “Stimulus subset profiles are inconclusive” denotes the neurons where the corresponding stimulus subsets activated only a small number of neurons (< 6), thus their responsiveness is inconclusive. (**H**) 15 ligand subsets that activated distinct neuronal subpopulations. It is obtained from the 32 stimulus subsets in (**F**) after merging stimulus subsets containing the positive controls (FF, MF) and different concentration conditions (1 µM, 10 µM).

Initially, we combined the marginal responsiveness for each VSN to determine a stimulus subset to which the VSN responded (each VSN’s chemosensory “tuning profile”). With this approach, we found that the simple combination of marginal responsiveness led to the identification of 443 distinct stimulus subsets activating at least one VSN (Figure 6D). But the majority of the detected stimulus subsets (307 out of 443) activated only one VSN among the 1,628 neurons. Given that there are only ∼300 known chemosensory receptors (VRs) expressed by this population [25, 26], and that the stimulus panel encompasses only a small fraction of the detectable chemosignals, it seemed highly unlikely that all the detected stimulus subsets reflected the underlying biology. This led us to consider many of these patterns as false positives. This potential discrepancy is an inherent limitation of the simple combination approach, in which the determination of marginal responsiveness by the FDR procedure carries a 10% chance of false positivity and the subsequent combination of these values introduced a higher chance of inaccuracies, especially when dealing with a large graph of ligand-neuron mapping.

To overcome the difficulty in controlling false positive combinations of stimulation-response pairings, we devised a selection approach based on a simple thresholding procedure. We first visualized the distribution of the number of VSNs activated by the identified stimulus subsets (Figure 6E). With an increasing number of activated VSNs in the distribution, the associated number of stimulus subsets decreased exponentially. This aligned with the typical distribution pattern observed in the occurrence of random events, specifically referring to the number of random false positivity events occurring for individual VSNs in this case. However, the distribution deviated from the exponential decay when the number of activated VSNs reached 6, displaying that the number of stimulus subsets that activated 6 VSNs exceeded the one that activated 5 VSNs (Figure 6E). Based on this observation, we concluded that stimulus subsets that activated more than 6 VSNs in this dataset were unlikely to occur by random chance. Therefore, we focused subsequent analyses on stimulus subsets that activated at least 6 VSNs.

This cutoff resulted in just 32 qualifying stimulus subsets (Figure 6F, See Methods). Of the 1,628 VSNs, 403 (25%) were determined to be unresponsive to any stimuli based on t-testing and FDR control, and 574 (35%) had response patterns where stimulus subset profiles were inconclusive because they did not exceed the inclusion threshold, resulting in a total of 651 VSNs (40%) with qualifying response patterns (Figure 6G). The most common response pattern was exclusive activation by male feces extract (MF), which was observed in 99/651 VSNs (15%). This was unsurprising; MF is natural extract containing many chemical ligands, and is therefore likely to activate multiple VRs (i.e., multiple populations of VSNs) [30]. The second most common response pattern was exclusive activation by 10 µM E0893 (91/651 VSNs, 14%), a sulfated estrogen known to activate multiple VRs at this concentration [31, 39]. The next most prevalent activity patterns included responses to both 10 µM E0893 and P3865 (65/651 VSNs, 10%), FF and MF (40/651 VSNs, 6%), etc. (Figure 6F). Qualifying response patterns also included exclusive activation by 10 µM CDCA (12/651 VSNs, 1.8%) and by both 1 µM CDCA and 10 µM CDCA (6/651 VSNs, 0.9%), because the analyses treated the different concentrations as different stimuli at this point. This is consistent with other studies suggesting VSNs express VRs that can be distinguished based on their ligand sensitivities [23, 31, 39]. These results indicate that by removing contributions from noise and false positivity our thresholding procedure produces biologically consistent results.

### DyNT identifies at least 15 subpopulations of VSNs that respond to distinct subsets of four bile acids and four sulfated steroids

To further simplify the analysis, we focused on just 8 stimuli (the 8 monomolecular ligands), collating VSN responses to the same ligand and different concentrations. VSNs activated by the stimulus subsets that included a stimulus FF or MF were merged into groups with the same response pattern as to monomolecular ligands. For example, ‘E0893 + P3865 + MF’ was merged into ‘E0893 + P3865’. Altogether, these actions converted 32 stimulus subset profiles into 15 ligand subset profiles (Figure 6H). Since VSNs predominantly express a single VR and VSNs with multiple VRs are scarce [40], neurons sharing a ligand subset profile (chemosensory tuning profile) are candidates to express the same VR or a small set of VRs that bind to every ligand in the profile but not to the other applied ligands. In other words, each of the 15 VSN subpopulations identified by their chemosensory profiles is likely to represent groups of neurons expressing independent VRs. Individual response examples of VSNs belonging to each of the 15 subpopulations are presented in Figure S2 and S3, displaying robustly identified chemosensory profiles of VSNs associated with the 8 applied bile acids and sulfated steroids.

Among the 15 subpopulations, 8 displayed exclusive responsiveness to one of the 8 monomolecular ligands (Figure 6H), supporting the presence of VRs that bind to only one ligand among the applied 8. The presence of such VRs may be useful for identifying specific biologically relevant molecules present in social odors. The remaining 7 neuronal subpopulations possessed broader chemosensory tuning profiles, including populations that responded to at least one bile acid and one sulfated steroid. This indicates the presence of VRs with broader ligand sensitivities, which may support a broad combinatorial basis upon which to build a representation of social odor identity [27, 30, 41].

Because the C = 6 threshold for inclusion of stimulus subset profiles may have limited these interpretations, we generated alternative profiles under the criteria that C = 3, 5, and 9 VSNs, which after merging resulted in 31, 17, and 12 ligand subsets, respectively (Figure S4). The C = 3 caused a massive expansion of tuning profiles, most of which were linear combinations of those observed when C = 6 (e.g., ‘CA + E0893’, ‘CDCA + E0893’, ‘E0893 + Q1570’, etc.). Given the acknowledged “crossover” effects caused by spatial overlap between nearby ROIs, it seems likely that these extra chemosensory profiles, which were observed only in a small number of neurons, were the result of signal contamination. In contrast, for the criteria C = 5 and C = 9 the activation profiles were similar to C = 6. Importantly, at C = 5, 6, and 9, the analysis indicated VSN subpopulations specific to each applied ligand. Cumulatively, these results indicate that our thresholding-based statistical procedure identifies robust combinatorial responsiveness profiles, revealing high-likelihood receptor-ligand profiles.

### Responsivity of VSNs to bile acids and sulfated steroids

Assuming most VSNs express a single VR, the number of VSN subpopulations activated by each applied ligand indicates the minimum number of VRs associated with a ligand. At the population level, there was a clear bias in the estimated number of associated VRs to two sulfated steroids, E0893 (6/15 VSN subpopulations) and P3865 (5/15 subpopulations; Figure 7A). This strong bias was partially expected, as E0893 (17α-estradiol sulfate) is a member of a well-studied chemosensory steroid family with multiple confirmed receptors [31, 33, 39]. Among the bile acid ligands, DCA contributed to the most tuning profiles (4/15). Two of the DCA-responsive profiles overlapped with P3865 and/or E0893. On the other hand, 10 µM Q1570 and 10 µM CDCA each participated only in a single, exclusive ligand subset profile (Figure 6H and Figure 7A). In addition, the identified ligand subset profiles also allowed us to estimate the minimum number of common VRs which are associated with each pair of ligands. For example, three of the 15 VSN subpopulations were activated by both E0893 and P3865, suggesting that at least three VRs bind to both (Figure 6H). Visualizing these estimates as a graph, one can appreciate that, as a population, VSNs combine information from ligands like E0893, P3865, and DCA with multiple other ligands in a way that over-represents these cues compared to others (Figure 7B).

**Fig. 7.**
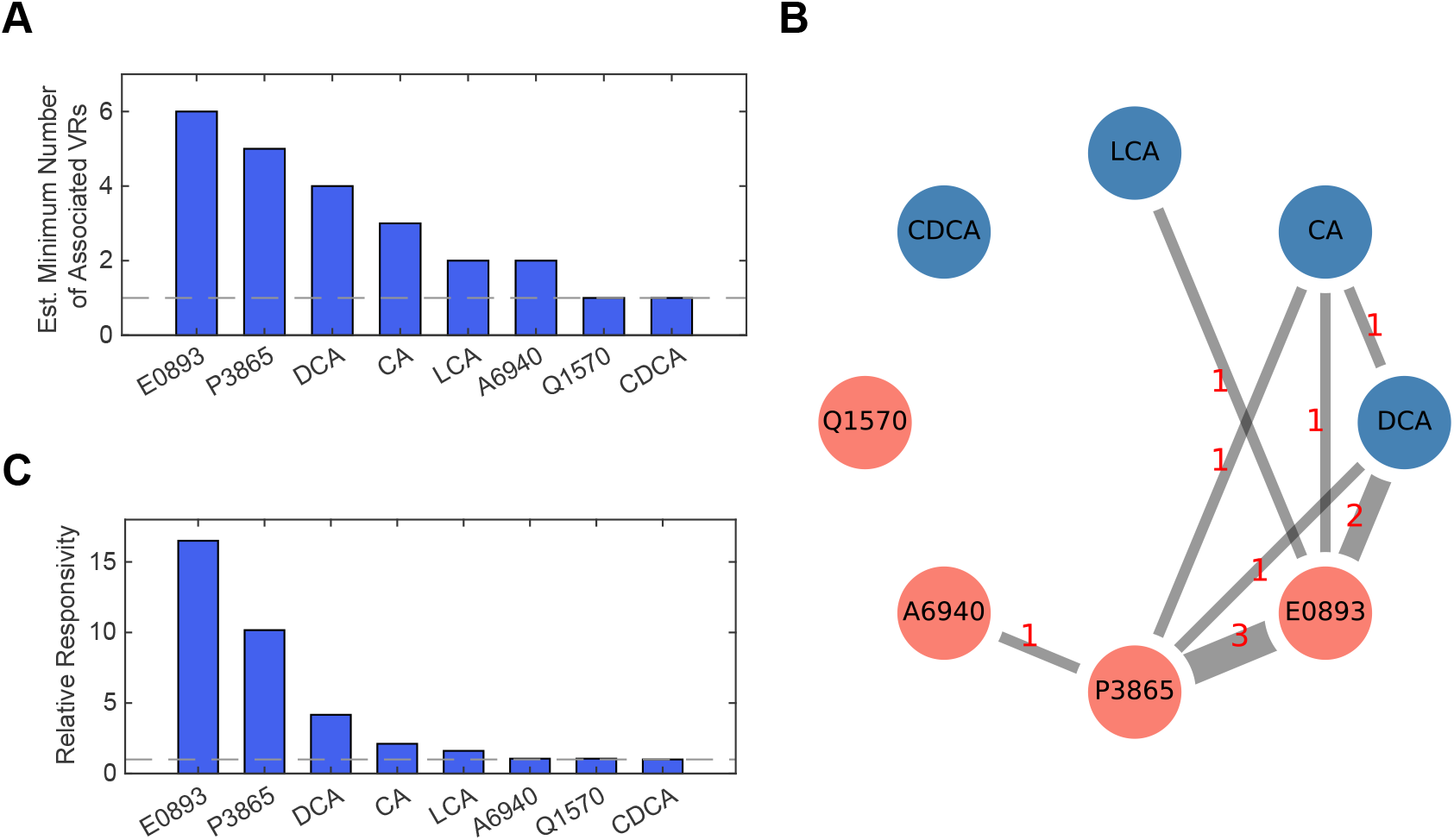
Responsivity of VSNs to applied four bile acids and four sulfated steroids. (**A**) The estimated minimum number of VRs associated with each ligand. The dotted gray line denotes the minimum number. (**B**) The estimated minimum number of common VRs shared by each pair of ligands (in red). Edge widths in the graph are proportional to the estimated numbers. (**C**) The relative responsivity of VSNs to the 8 applied ligands. The number of VSNs activated by each ligand is normalized to the smallest observed count, which is 18 for CDCA. The dotted gray line denotes the minimum of one.

The over-representation of certain ligands across tuning profiles was paralleled by over-representation in the total number of neurons activated per ligand. For example, 10 µM E0893 activated the largest number of VSNs (297), while 10 µM CDCA activated the smallest number of VSNs (18). We normalized the number of activated VSNs per ligand by the smallest count to quantify the relative responsivity of VSNs to the applied 8 ligands (Figure 7C). The largest relative responsivity was 16.5 for E0893, followed by P3865 (10.2) and DCA (4.2). These relative responsivities mirrored the order of the estimated minimum numbers of associated VRs (Figures 7A and 7C). However, the gradient of this disproportionate responsivity was steeper than the gradient in the estimated numbers of associated VRs. For example, E0893 activated approximately 16.5 times more VSNs than CDCA, while it was likely to possess about 6 times more associated VRs than CDCA. This suggests that the expression of VRs associated with E0893 or P3865 in VNO tissues may be higher than that of Q1570 or CDCA. Overall, these data indicate that, when concentrations are matched, VSNs have a noted bias towards specific molecules, and that the cellular mechanisms underlying this phenomenon are worthy of further investigation.

## Discussion

Calcium fluorescence imaging has greatly advanced our understanding of neural circuits in living animals. At the same time, the technology has raised significant computational challenges in processing and interpreting massive amounts of neuronal activity data it generates. A key challenge lies in the registration of brain tissue undergoing non-linear and erratic deformations during prolonged recording periods. This registration requires single-cell precision throughout the entire duration of the movie, which often cannot be accomplished by existing calcium imaging pipelines relying on course-grained registration and/or fixed ROI segmentation.

In this study, we present DyNT as a tool to generate dynamic ROIs encompassing highly active individual neurons in deforming tissues. While developing the proposed method, we prioritized accuracy, because any mapping of ligand-receptor association strongly depends on the consistency of ROIs capturing single neurons. A key algorithmic component of DyNT is the patch-matching iteration step, where neuron trajectories and their local mean images update each other iteratively. This iteration was motivated by the expectation-maximization algorithm [42]. It searches for the best connections between neuronal firing events at different time points that are supposed to originate from one neuron.

Motion artifacts (both transient and sustained) disrupt not only the neuron segmentation, but also the confidence in attributing calcium changes to individual neurons in each ROI. Given that many calcium imaging experiments involve highly active, independent, adjacent neurons, the ability to minimize crosstalk in the readout of calcium activation is critical [8, 19]. Several studies previously reported overlaps of adjacent neurons in two-photon calcium imaging data due to the projection of a shallow volume to a 2D plane [15, 19]. To mitigate the signal interference between neighboring neuronal activities, the DyNT pipeline incorporates a weighting scheme that utilizes the information from neuronal mean images and spatial correlations. This approach diminishes spurious activity peaks in many neuronal activity time courses (Figure 3C and 3D). It will be useful for enhancing the accuracy of measuring single neurons’ calcium activities in general calcium imaging data, regardless of whether the activity measurements are based on fixed or dynamic ROIs.

A key current limitation of DyNT is a long execution time. For example, it took ∼330-560 min to process individual sub-volume VNO videos with file sizes of ∼1 GB, using a high-performance computing node with 32 CPUs and 384 GB RAM. This contrasts to CaImAn’s processing time of ∼4-8 min for the same data sets. However, it should be noted that CaImAn’s fixed ROI method generated a significant portion of incorrect ROIs, which could not be removed or corrected by simply adjusting control parameters. To accurately track the positional jitters, DyNT relies on several iterative optimization procedures that are inherently costly. To accelerate computation in future versions, it will be necessary to translate high-level codes into more efficient, machine-proximal codes.

The DyNT pipeline, like others [8, 9, 12, 15, 19] produces ROIs without extensive user interaction, and without the need to incorporate stimulus identity or timing information into the segmentation workflow. This allows the identification and analysis of spontaneously active neurons, which are difficult to segment manually. If calcium imaging experiments are implemented with a set of shuffled stimulus repeats, the DyNT pipeline additionally produces statistical outputs about the combinatorial responsiveness of neurons. Integrated statistical analyses of marginal responsiveness determine which stimulus patterns are unlikely the result of biological noise or spurious overlap between adjacent ROIs. Using these features of DyNT, we studied combinatorial coding in the context of pheromone-sensing in the mouse VNO, a process that involves the detection and discrimination of complex blends of excreted chemicals. Using a targeted panel of 8 well-described ligands, including 4 bile acids and 4 sulfated steroids, we identified 15 VSN subpopulations with distinct ligand tuning patterns. These tuning patterns predict that, at the common 10 µM concentration, there exist at least 15 VRs sensitive to this panel. 8 of the 15 subpopulations were exclusively activated by a single ligand, while 7 responded to multiple ligands.

The combinatorial nature of the observed chemosensory tuning profiles supports the hypothesis that VRs, as a family, include receptors that have high sensitivity and selectivity for specific ligands, as well as receptors with broad ligand responsivity, potentially including receptors that sense multiple ligand classes [23, 27, 30, 39, 43, 44]. It is intriguing that the response profiles we identified markedly differ from those expected under a hypothetical strategy where different VRs would be randomly assigned to one or multiple ligands among the applied 8. In this study, E0893 (17α-estradiol sulfate) contributed the most tuning profiles (6 profiles; Figure 6H, Figure 7A). E0893 is an endogenous steroidal estrogen derivative, and a member of a class of estrogens with high chemosensory potency [31, 33, 39]. Sulfated estrogens, as a class, were previously shown to activate many VRs [31, 33], but these data raise several questions regarding the relationship between VR expression and VSN tuning.

Namely:

(A) Do bile acid-sensitive VRs have a single ligand binding site that is sensitive to both bile acids and estrogens (and vice-versa)?
(B) Do bile acid-sensitive VRs have multiple ligand binding sites, one for bile acids and one for estrogens (and vice-versa)?
(C) Do some VSNs express multiple receptors, including bile acid-sensitive and estrogen-sensitive receptors? If so, are estrogen-sensitive VRs commonly co-expressed with other VRs?

Studying these topics will require targeted future studies, but it is important to note that DyNT revealed these features using automated segmentation and statistically based stimulus tuning evaluation. This makes the approach especially attractive as a platform for studying combinatorial coding in chemosensory systems, where high-dimensional datasets are increasingly used because of the complex nature of this sensory modality [45].

## Materials and Methods

### Experimental Design

This study proposes a neuron segmentation method and software for 3D calcium imaging data. Most of the existing computational pipelines for single neuron segmentation generate fixed ROIs assuming that raw imaging data has been well-registered and neuron locations are static in the registered calcium videos. In contrast, our pipeline is designed for the case when calcium image registration is imperfect, mainly due to complicated tissue deformation, so neuron positions are dynamic over time. The DyNT pipeline written in MATLAB generates dynamic ROIs for moving single neurons and extracts accurate calcium activity time courses from 3D calcium imaging videos. In addition to neuron segmentation, the DyNT pipeline includes a statistical module to determine the combinatorial responsiveness of sensory neurons when neuronal activities have been recorded while exposed to repeated stimulus delivery.

### Imaging experiments and preprocessing

We analyzed volumetric calcium imaging data of acutely dissected mouse VNOs [23]. The VNOs were dissected from Omp-cre mice (Omp^tm4(cre)Mom^/MomJ) [46] crossed to “Ai96” mice (Gt(ROSA)26Sor^tm96(CAG-GCaMP6s)Hze^/J) [47], which express the genetically encoded calcium indicator GCaMP6s in chemosensory neurons. VNOs were imaged by using 3D light-sheet microscopy (objective-coupled planar illumination, OCPI) [48] at the frequency of 3 sec/frame (0.33 Hz). For the entirety of imaging sessions, Ringer’s saline solution was superfused over the tissue at a rate of ∼1 mL/min. To investigate ligand-VR interactions, the following monomolecular ligands were dissolved into Ringer’s solution: the bile acids CA, DCA, LCA, and CDCA (each at 1 µM and 10 µM) and the sulfated steroids A6940, E0893, P3865, and Q1570 (each at 10 µM). The four bile acids and four sulfated steroids were delivered every 60 sec in a randomized order, repeated multiple times. Each stimulus presentation lasted for 15 sec followed by a rest period of 45 sec. The raw imaging data were down-sampled for computational efficiency, and rigid-registered using phase correlations. In the down-sampled image volume, the spatial resolution was 2.8 µm/pixel in the X/Y-axis and 8 µm/pixel in the Z-axis.

### Multi-scale detection of firing neurons and tracking small jitters

The first step of the DyNT pipeline is to detect firing neurons and track their movements within consecutive time frames. For particle detection, DyNT utilizes an algorithm based on Laplacian-of-Gaussian filtering implemented in u-track 3D [36]. As described for 2D images in our previous study [49], the detection algorithm searches for candidate particle locations by examining the local fitness of a 3D Gaussian point spread function. Hence, the standard deviation parameters (σ_X_, σ_Y_ and σ_Z_) determine the optimal size and shape of particles to be detected. The DyNT pipeline takes multiple choices for the standard deviation parameters of 3D Gaussian functions and detects firing neurons of various sizes and shapes. In processing the VNO videos in this paper, we used three choices of standard deviation parameters to capture different sizes ([σ_X_=σ_Y_, σ_Z_]∈{[9.8, 12], [5.6, 8], [2.8, 8]} µm). The pipeline is designed to process any number of different scales, which depends on individual imaging data.

After the detection of firing neurons at each time frame, DyNT identifies firing events by tracking small movements of detected firing neurons within consecutive time frames. Because the small spatial jitters observed between some image stacks are simple in terms of particle dynamics, the DyNT pipeline provides a pre-specified tracking parameter object (*TrackingParams_init.mat*) which is fed into the u-track 3D package [36]. The tracking parameters specify simple particle dynamics with no directed motion, minimal gap closing (1 frame), 1-2 pixels of Brownian motion search radius, etc. Hence, DyNT requires no user parameters for tracking small, transient displacement.

### Tracking dynamic neurons using multiple correlation thresholds

A large number of the detected firing events are fed into patch-matching iterations to identify which firing events at different time frames belong to identical neurons based on spatial correlations. To compute local mean images of firing events, DyNT requires user input for local patch size (Figure 2C). Users also need to specify maximal jittering amounts in X/Y-axis and Z-axis, so that a local patch for a single neuron is assumed to move around over time within a neighborhood volume that is wider by 2 × (maximal jittering amounts) in each axis.

Rolling spatial correlations between the local patch and its neighborhood volume can help pinpoint when and where the corresponding firing neuron appears again. DyNT exploits the maximum of rolling spatial correlations at each time frame and determines the corresponding neuron to be firing if the maximum correlation is greater than a pre-specified threshold (Figure 2E). We found that using multiple thresholds for the correlations substantially increased the number of segmented or tracked neurons. The DyNT pipeline runs two sets of patch-matching iterations with the different thresholds of {0.9, 0.8} and then merges the converged neuron trajectories after assessing if each pair of trajectories is essentially the same or not. It seems that the lower threshold of 0.8 works better for the case when neurons are densely populated, and adjacent neurons are co-firing in the local volume patch. Hence, locally heterogeneous neuronal activities require different optimal correlation thresholds, and using multiple thresholds leads to better segmentation performance.

### Moving median normalization and a weighting scheme for calcium activity measurement

Once dynamic ROIs are obtained, the averaged fluorescence intensities within the ROIs across the time frames generate raw calcium activity time courses for segmented single neurons. For visualization and subsequent analysis of activities, we normalize the raw calcium activities by using their moving medians as the baseline activities. The moving median of the raw activities {*I*(*t*)} of a single neuron at time *t* is denoted by *movMedian*(*I*(*t*), *w*), where *w* indicates the size of moving windows in frames within which a median of the activities is computed. Then, the moving median normalized calcium activities are given by:

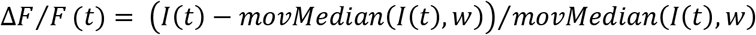

The DyNT pipeline further processes the normalized calcium activities to reduce spurious peaks and noise induced by adjacent neuronal activities (Figure 3). Because the maxima of rolling spatial correlations after iterations converge are indicative of whether the target neuron is active or not, we utilize the maximum spatial correlations, {*maxSpatialCorr*(*t*)}, after truncation as multiplicative weights:

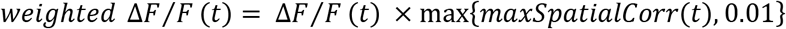

### Determining marginal responsiveness via t-tests and an FDR controlling procedure

The statistical module in the DyNT pipeline begins with determining the marginal responsiveness of segmented neurons to a given set of stimuli. In the VNO videos, a total of 15 stimuli were applied including 8 monomolecular ligands at different concentrations, two positive controls and one negative control (Ringer’s). Each of 15 different stimuli was applied 5 times during the 75-minute recording. Each trial period of 20 frames (60 sec) was chopped and calcium activity response curves across trials for each neuron and stimulus were collected. The activities of the first 10 frames within each trial were averaged, resulting in a simple dataset of five average response values for each neuron and each stimulus. For each neuron and stimulus except Ringer’s, the five response values were compared with the responses for Ringer’s using a one-sided t-test. Since it is a multiple-comparison situation, the P-values were adjusted using a Benjamini-Hochberg FDR controlling procedure [37].

### Stimulus delivery information input and combinatorial responsiveness profile outcome

The statistical module in the DyNT pipeline automates statistical analyses for calcium imaging data to address the overarching question of how sensory neurons encode a given set of environmental inputs. DyNT assumes imaging experiment setups where multiple stimuli are applied sequentially in a randomized order, multiple times, and at every fixed time interval. If the stimulus delivery information is provided, the statistical module determines the combinatorial responsiveness of each segmented neuron to an applied set of stimuli based on multiple calcium video data. The stimulus delivery information can be provided by a *.csv* file for each video, which needs to contain three columns with the information of ‘*stimulusLabel*’, ‘*startFrame*’, and ‘*endFrame*’.

All the segmented sensory neurons from multiple videos are determined into one of three categories (Figure 6G): (i) non-responsive to any given stimulus; (ii) the case where the responsiveness profile is inconclusive because the corresponding stimulus subset activates only a small number of neurons; (iii) the case where a stimulus subset is identified for the neuron. For the third case, the identified stimulus subsets and the number of neurons activated by the subsets are presented through dot visualization (Figure 6F). Furthermore, the DyNT pipeline generates all the relevant information such as positions of neurons, local MIP videos of the segmented neurons, across-trial response curve plots (Figure 5), and boxplots for the t-tests and FDR control, all of which are annotated with the determined responsiveness profiles so that users can easily locate and characterize neurons of particular interest.

### Statistical analysis

To determine the marginal responsiveness of individual VSNs, we used one-sided t-tests to test if the first 10-frame averaged calcium responses to a stimulus were greater than the responses to Ringer’s (Figures 5-6). Then the P-values computed over multiple stimuli and VSNs were adjusted using a Benjamini-Hochberg FDR controlling procedure [37].

## Acknowledgments

Technical support for imaging experiments was provided by Cara Nielson. We thank members of the Meeks and Danuser laboratories for advice and feedback.

## Author contributions

WMW and JPM designed the imaging experiments. WMW performed experiments and performed manual ROI annotations. JN and JPM developed the DyNT objectives. JN developed DyNT software and subsequent analysis. JN, JPM, and GD interpreted results and wrote the paper.

## Data and materials availability

All data are available in the main text or the supplementary materials. Data for the figures presented here are available in a public database (DOI: 10.17632/ny7wnxz8ry.1). All original codes for DyNT including example datasets, documentation, and a demo script are publicly available (https://github.com/JungsikNoh/DynamicNeuronTracker).

## Supplemental Figure Legends

**Fig. S1.**
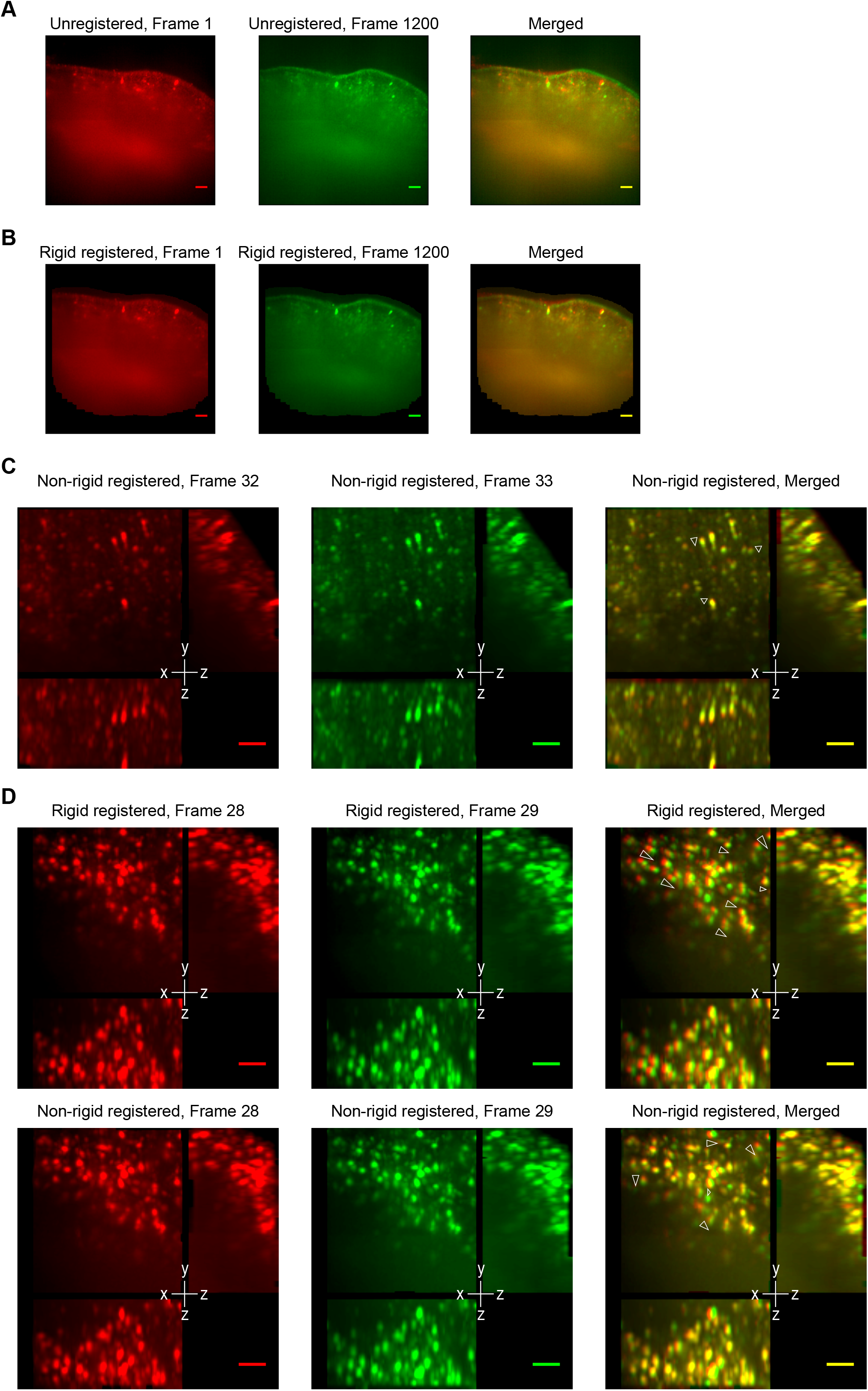
Jittering neurons in 3D calcium imaging of deforming tissues. (**A**) xy-plane images (the 15^th^ z-stack) of a VNO unregistered video at the first and last frames. The tissue boundaries and constantly active neurons display mild tissue deformation during an hour of imaging. All scale bars, 50 µm. (**B**) xy-plane images of the VNO video in (**A**) after rigid registration. The deformation is reduced but remains. (**C**) After the sub-volume video for Figure 1A is non-rigid registered, jittering neurons are annotated (white triangle) on MIP images at the same time frames in Figure 1A. (**D**) After another VNO video is rigid (top) and non-rigid (bottom) registered, jittering neurons are annotated on consecutive MIP images.

**Fig. S2.**
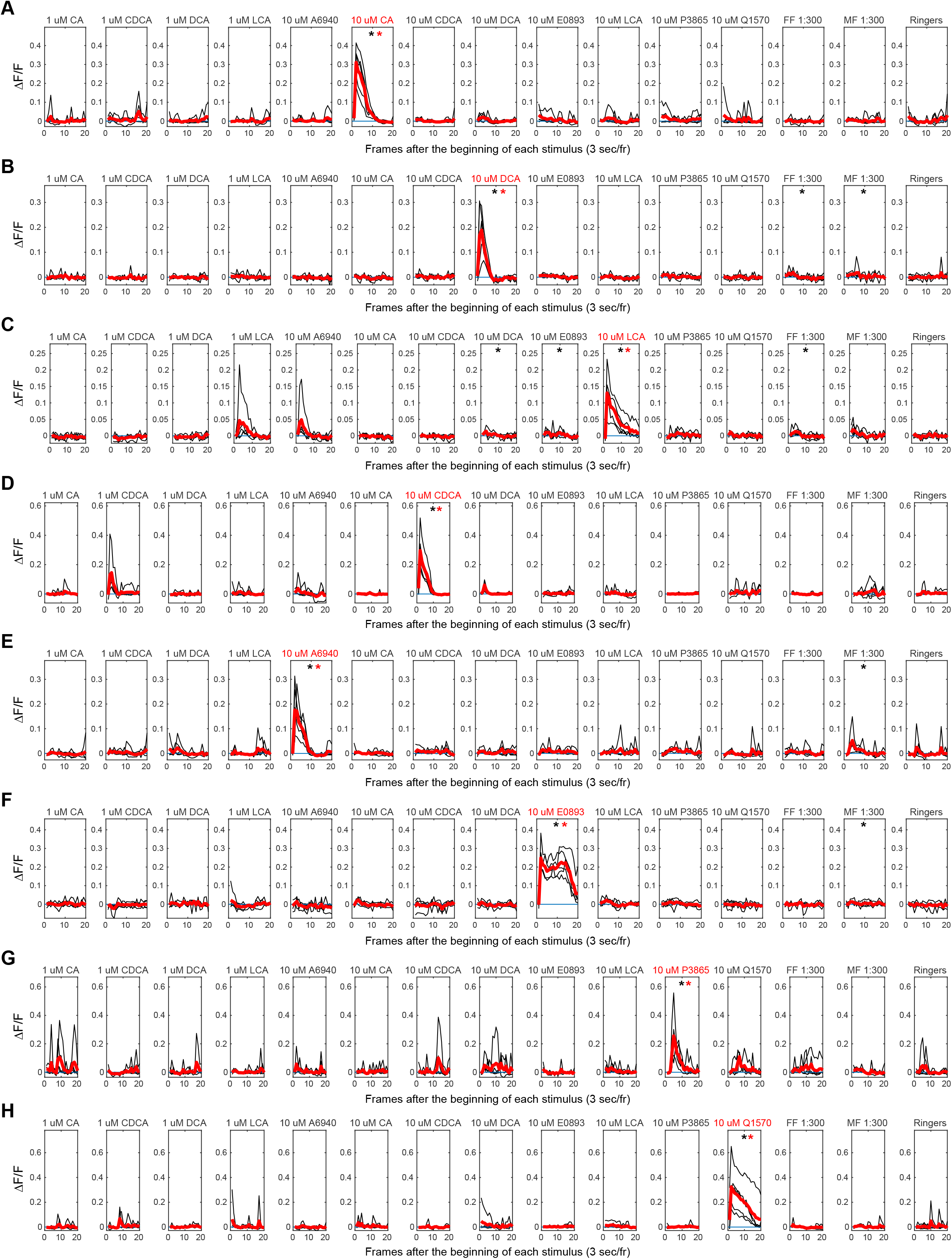
Across-trial responses of representative 8 VSNs that exclusively respond to one of applied four bile acids and four sulfated steroids. (**A**-**H**) Calcium activity response curves (black) of the representative 8 VSNs that exclusively respond to CA, DCA, LCA, CDCA, A6940, E0893, P3865, or Q1570, respectively. The red curve indicates the averaged response curve over five trials for each stimulus. The black star indicates that the across-trial responses for a stimulus are significantly higher than the ones for Ringer’s (P < 0.05, one-sided t-test). The red star indicates significance after FDR control (Adjusted P < 0.1).

**Fig. S3.**
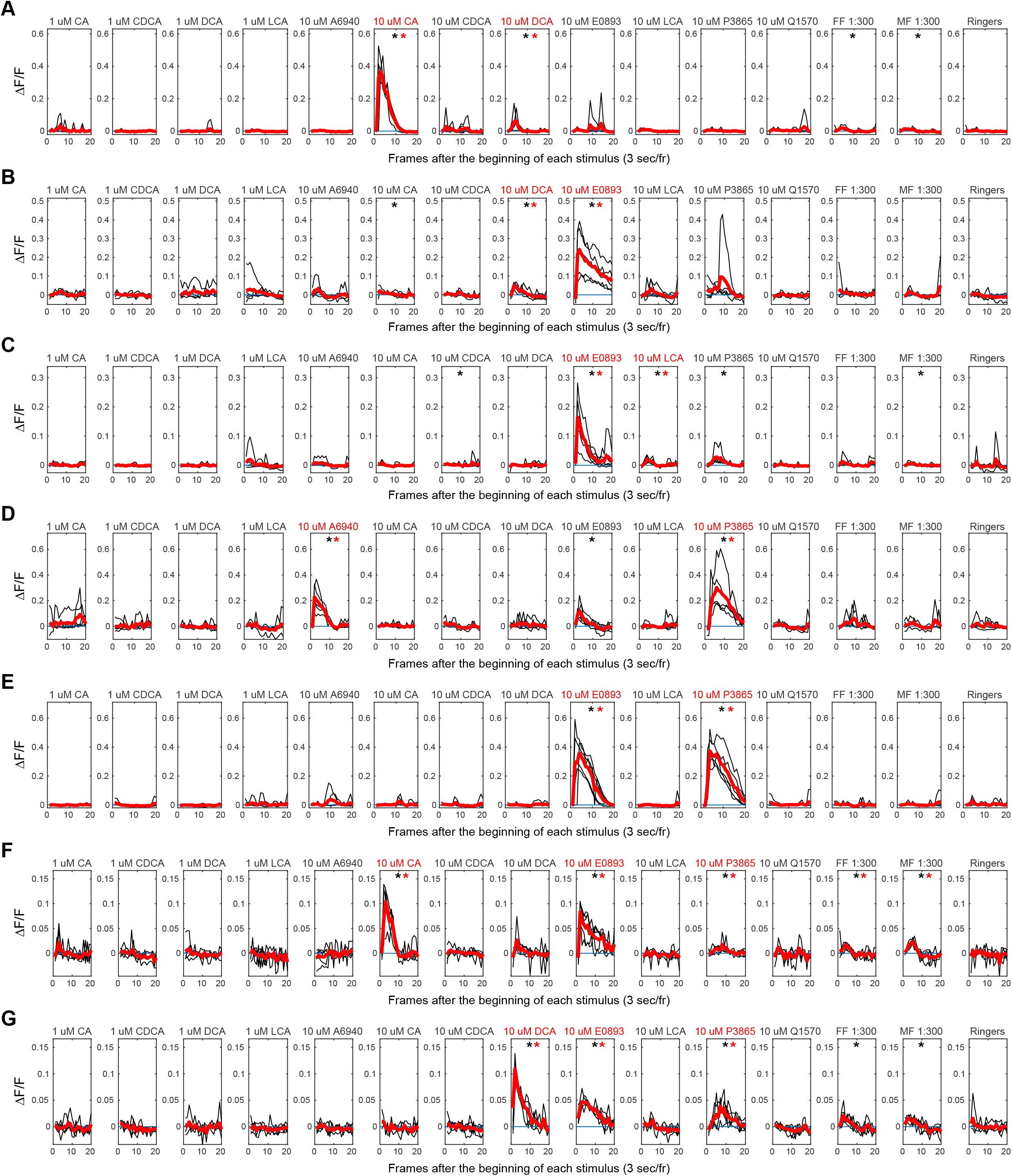
Across-trial responses of representative 7 VSNs that exclusively respond to distinct subsets of applied 8 ligands. (**A**-**G**) Calcium activity response curves (black) of the representative 7 VSNs that exclusively respond to {CA, DCA}, {DCA, E0893}, {E0893, LCA}, {A6940, P3865}, {E0893, P3865}, {CA, E0893, P3865}, or {DCA, E0893, P3865}, respectively. The red curve indicates the averaged response curve over five trials for each stimulus. The black star indicates that the across-trial responses for a stimulus are significantly higher than the ones for Ringer’s (P < 0.05, one-sided t-test). The red star indicates significance after FDR control (Adjusted P < 0.1).

**Fig. S4.**
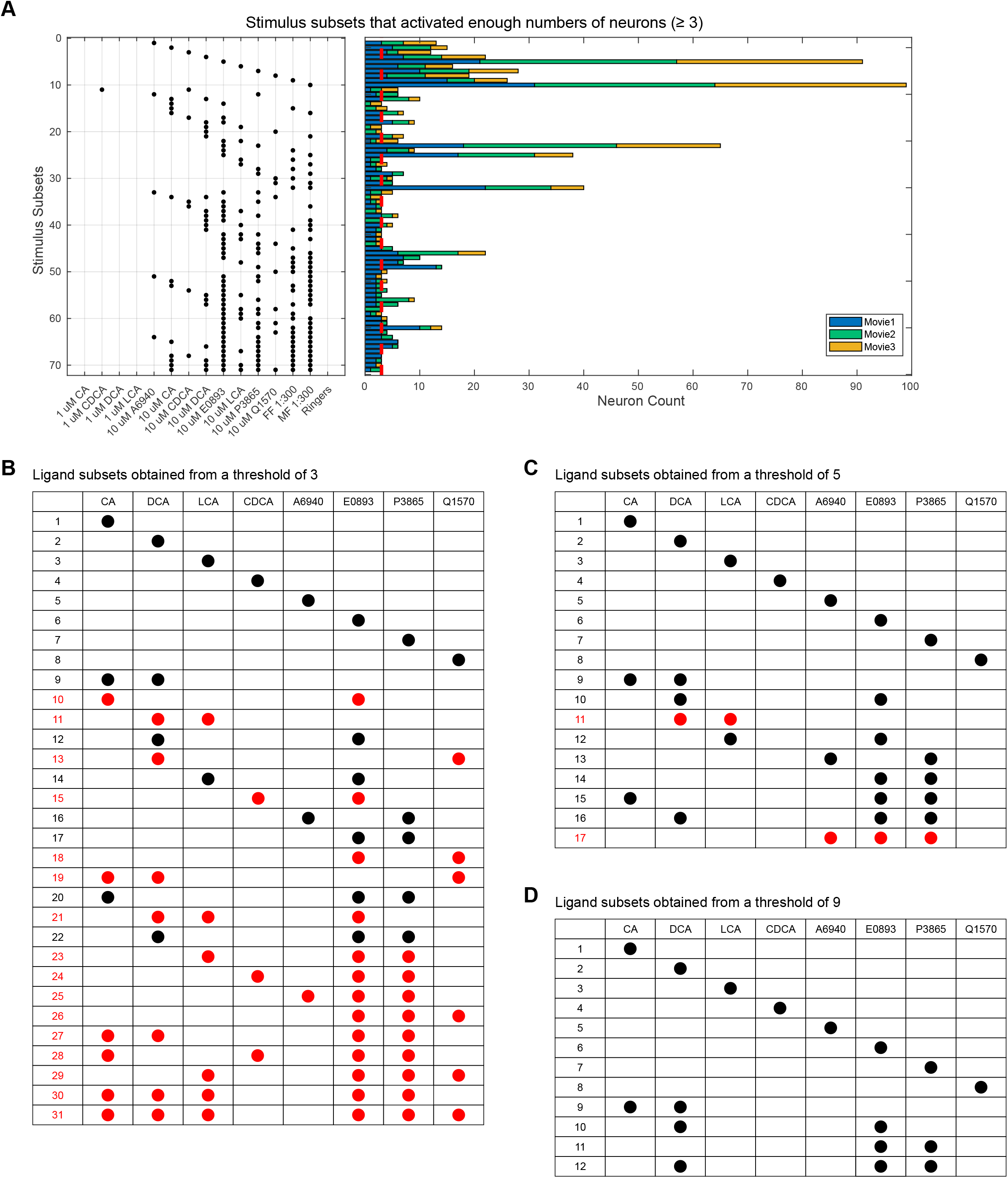
Combinatorial responsiveness profiles with different inclusion thresholds. (**A**) 71 stimulus subsets are identified when we apply a threshold of three, that is, after selecting the stimulus subsets that activated at least three VSNs. Movie indexes of the activated neurons are color-coded. (**B**) 31 ligand subsets are derived from the 71 stimulus subsets in (**A**) after merging the positive controls (FF, MF) and different concentration conditions (1 µM, 10 µM). Rows with black dots indicate the same ligand subsets identified by the threshold of six as in Figure 6H. Rows with red dots indicate additional ligand subsets identified by the lower threshold of three. (**C**-**D**) 17 (12) ligand subsets are identified when a threshold of five (nine) is applied, respectively.

## Supplemental Video Legends

**Video S1. A xy-plane of an unregistered 3D calcium video of VNO tissue displays mild non-linear tissue deformation during an hour of imaging, related to Figure S1.**

The xy-plane is the 12^th^ z-stack among 51 stacks. The imaging time interval is 3 sec/frame. Fast replay at 100 frames/second. Scale bar, 50 µm.

**Video S2. Maximum intensity projection (MIP) of an unregistered 3D calcium video of VNO issue displays positional jitters of single neurons, related to Figure S1.**

MIP of the same calcium imaging data as Video S1. Fast replay at 100 frames/second. Scale bar, 50 µm.

**Video S3. Rigid (left) and non-rigid (right) registration of a sub-volume video of VNO tissue, related to Figure 1**.

Maximum intensity projection of a sub-volume video of VNO tissue after registrations. The single time frames of this video after rigid registration are shown in Figure 1. The sub-volume covers ∼7% of the full volume containing the tissue. Replay at 20 frames/second. Scale bar, 50 µm.

**Video S4. Rigid (left) and non-rigid (right) registration of a sub-volume video of VNO tissue, related to Figure S1D.**

Maximum intensity projection of a sub-volume video of VNO tissue after registrations. The single time frames of this video are shown in Figure S1D. Replay at 20 frames/second. Scale bar, 50 µm.

**Video S5. CaImAn (left) and DyNT (right) segmentation outcomes for a sub-volume video of VNO tissue, related to Figure 1**.

ROIs that contain high calcium activities are overlaid on MIP images. Different ROIs are color-coded. Replay at 20 frames/second. Scale bar, 50 µm.

**Video S6. CaImAn (left) and DyNT (right) segmentation of an example neuron (N1), related to Figure 1**.

A MIP video of a local volume around an example neuron (N1) in Figure 1 is shown. ROIs are displayed only when their calcium activities are high. The neuron is annotated by two CaImAn ROIs (red and yellow, left), and a single DyNT ROI (red, right). Replay at 20 frames/second. Scale bar, 50 µm.

**Video S7. CaImAn (left) and DyNT (right) segmentation of an example neuron (N2), related to Figure 1**.

A MIP video of a local volume around an example neuron (N2) in Figure 1 is shown. ROIs are displayed only when their calcium activities are high. The neuron is annotated by two CaImAn ROIs (yellow and orange, left), and a single DyNT ROI (yellow, right). Replay at 20 frames/second. Scale bar, 50 µm.

**Video S8. CaImAn (left) and DyNT (right) segmentation of an example neuron (N3), related to Figure 1**.

A MIP video of a local volume around an example neuron (N3) in Figure 1 is shown. ROIs are displayed only when their calcium activities are high. Its ROI by CaImAn (yellow, left) contains multiple neurons, but the ROI by DyNT (yellow, right) separates N3 from adjacent neurons. Replay at 20 frames/second. Scale bar, 50 µm.

**Video S9. CaImAn (left) and DyNT (right) segmentation of an example neuron (N4), related to Figure 1**.

A MIP video of a local volume around an example neuron (N4) in Figure 1 is shown. ROIs are displayed only when their calcium activities are high. Its ROI by CaImAn (yellow, left) contains multiple neurons, but the ROI by DyNT (yellow, right) separates N4 from adjacent neurons. Replay at 20 frames/second. Scale bar, 50 µm.

**Video S10. Dynamic ROIs generated by DyNT from a 3D calcium video of entire VNO tissue, related to Figure 3**.

Segmented 561 dynamic ROIs are shown when they are active on top of MIP images of an entire VNO tissue. Different ROIs are numbered and color-coded. The stimulation status during imaging is indicated at the lower right. Replay at 20 frames/second. Scale bar, 50 µm.

**Video S11. A MIP video of a local volume around a representative VSN that responds to only CA, related to Figure 5**.

The entire 1,500 frames are played at 20 frames/second. Neuron masks are shown only when they are active. The representative neuron is annotated in yellow at the center. The stimulation status is indicated at the lower right. Scale bar, 50 µm.

**Video S12. A MIP video of a local volume around a representative VSN that responds to only DCA, related to Figure 5**.

The entire 1,500 frames are played at 20 frames/second. Neuron masks are shown only when they are active. The representative neuron is annotated in yellow at the center. The stimulation status is indicated at the lower right. Scale bar, 50 µm.

**Video S13. A MIP video of a local volume around a representative VSN that responds to only E0893 and P3865, related to Figure 5**.

The entire 1,500 frames are played at 20 frames/second. Neuron masks are shown only when they are active. The representative neuron is annotated in yellow at the center. The stimulation status is indicated at the lower right. Scale bar, 50 µm.

**Video S14. A MIP video of a local volume around a representative VSN that displays activities non-specific to applied ligands, related to Figure 5**.

The entire 1,500 frames are played at 20 frames/second. Neuron masks are shown only when they are active. The representative neuron is annotated in yellow at the center. The stimulation status is indicated at the lower right. Scale bar, 50 µm.

