## Supplemental Figures 1-4 for "Combinatorial responsiveness of single chemosensory neurons to external stimulation of mouse explants revealed by DynamicNeuronTracker"

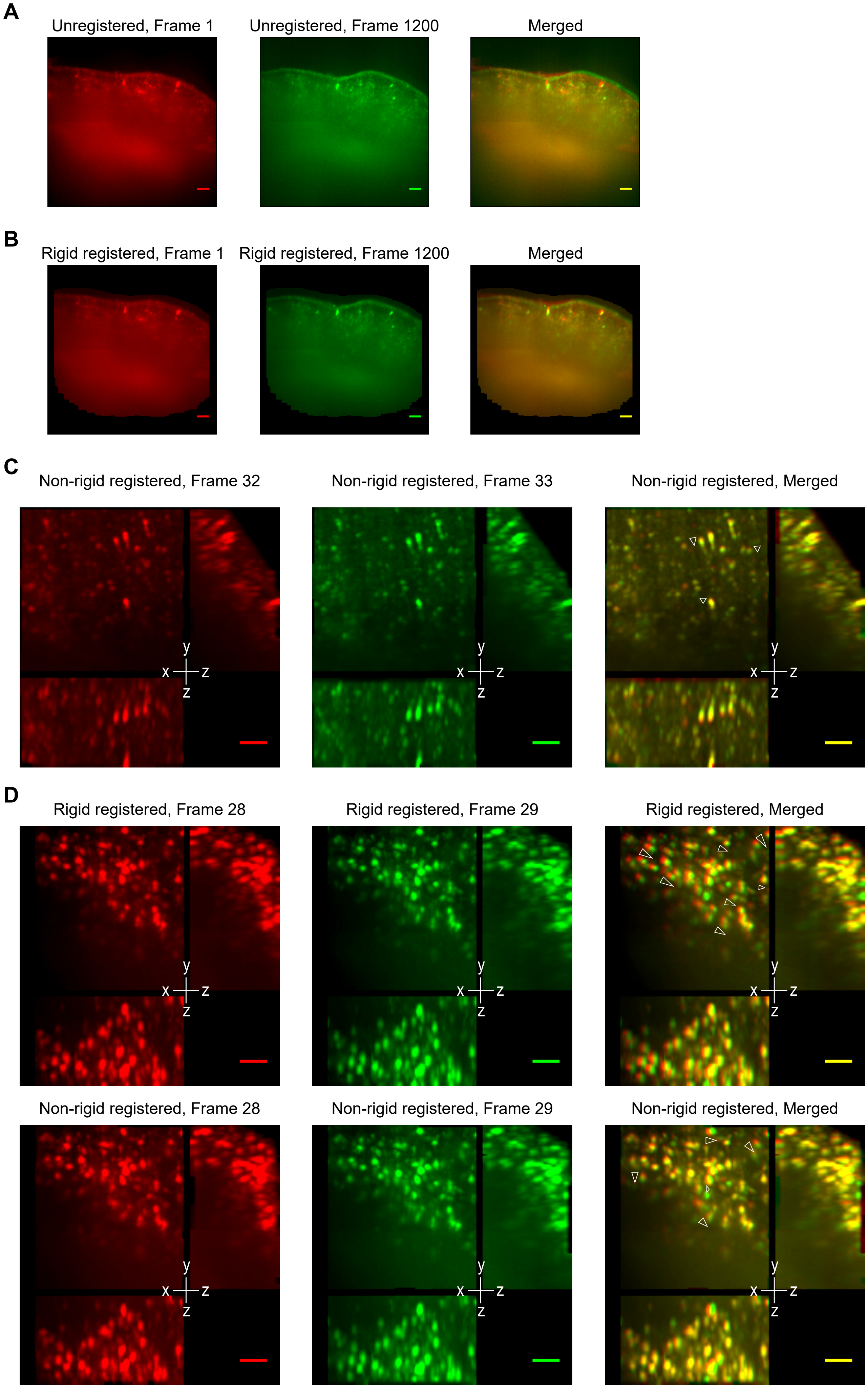
A
Unregistered, Frame 1
Unregistered, Frame 1200
Merged
B
Rigid registered, Frame 1
Rigid registered, Frame 1200
Merged
C
Non-rigid registered, Frame 32
Non-rigid registered, Frame 33
Non-rigid registered, Merged
D
Rigid registered, Frame 28
Rigid registered, Frame 29
Rigid registered, Merged
Non-rigid registered, Frame 28
Non-rigid registered, Frame 29
Non-rigid registered, Merged
y
x
z
z

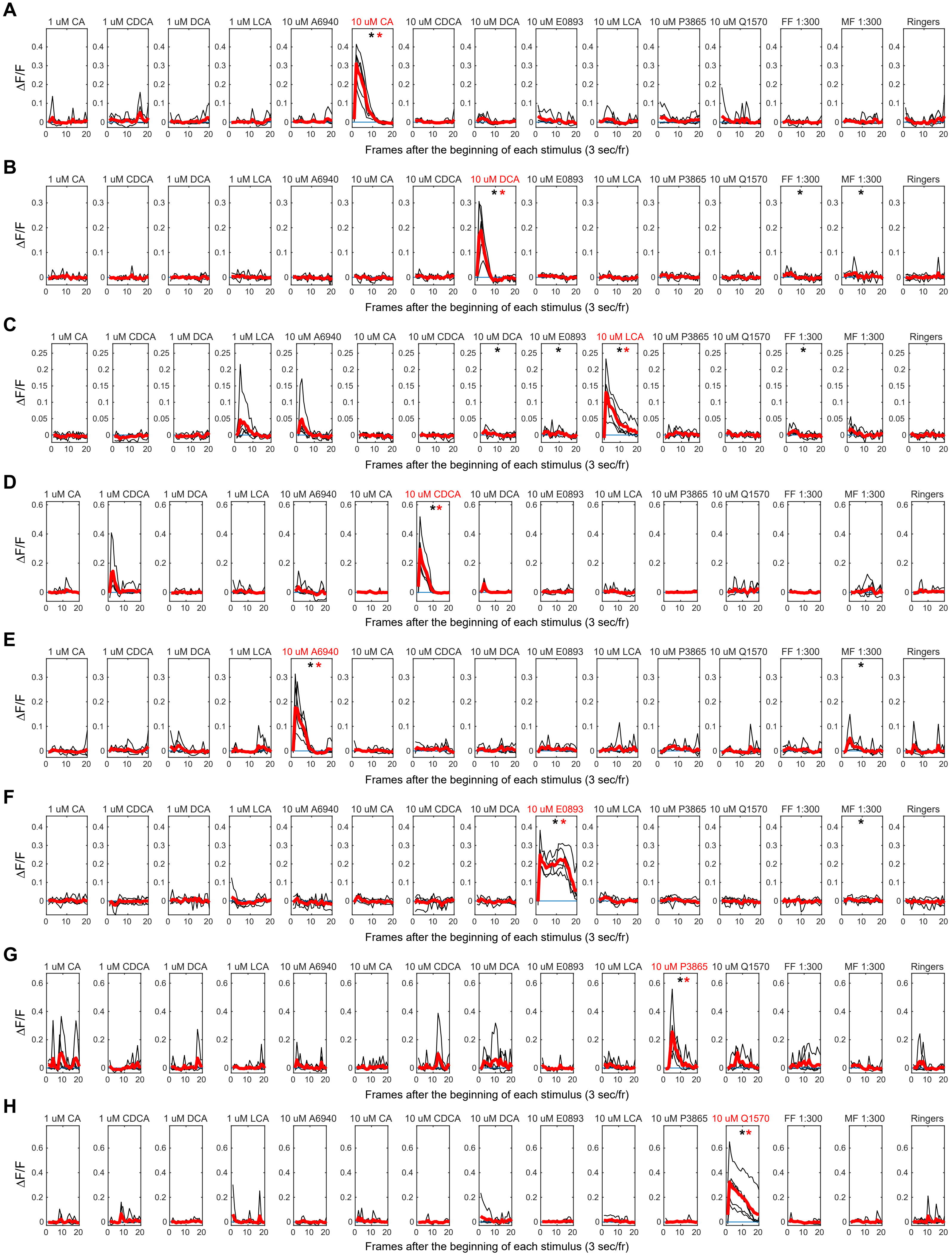
A
1 uM CA
1 uM CDCA
1 uM DCA
1 uM LCA
10 uM A6940
10 uM CA
10 uM CDCA
10 uM DCA
10 uM E0893
10 uM LCA
10 uM P3865
10 uM Q1570
FF 1:300
MF 1:300
Ringers
ΔF/F
Frames after the beginning of each stimulus (3 sec/fr)
B
1 uM CA
1 uM CDCA
1 uM DCA
1 uM LCA
10 uM A6940
10 uM CA
10 uM CDCA
10 uM DCA
10 uM E0893
10 uM LCA
10 uM P3865
10 uM Q1570
FF 1:300
MF 1:300
Ringers
ΔF/F
Frames after the beginning of each stimulus (3 sec/fr)
C
1 uM CA
1 uM CDCA
1 uM DCA
1 uM LCA
10 uM A6940
10 uM CA
10 uM CDCA
10 uM DCA
10 uM E0893
10 uM LCA
10 uM P3865
10 uM Q1570
FF 1:300
MF 1:300
Ringers
ΔF/F
Frames after the beginning of each stimulus (3 sec/fr)
D
1 uM CA
1 uM CDCA
1 uM DCA
1 uM LCA
10 uM A6940
10 uM CA
10 uM CDCA
10 uM DCA
10 uM E0893
10 uM LCA
10 uM P3865
10 uM Q1570
FF 1:300
MF 1:300
Ringers
ΔF/F
Frames after the beginning of each stimulus (3 sec/fr)
E
1 uM CA
1 uM CDCA
1 uM DCA
1 uM LCA
10 uM A6940
10 uM CA
10 uM CDCA
10 uM DCA
10 uM E0893
10 uM LCA
10 uM P3865
10 uM Q1570
FF 1:300
MF 1:300
Ringers
ΔF/F
Frames after the beginning of each stimulus (3 sec/fr)
F
1 uM CA
1 uM CDCA
1 uM DCA
1 uM LCA
10 uM A6940
10 uM CA
10 uM CDCA
10 uM DCA
10 uM E0893
10 uM LCA
10 uM P3865
10 uM Q1570
FF 1:300
MF 1:300
Ringers
ΔF/F
Frames after the beginning of each stimulus (3 sec/fr)
G
1 uM CA
1 uM CDCA
1 uM DCA
1 uM LCA
10 uM A6940
10 uM CA
10 uM CDCA
10 uM DCA
10 uM E0893
10 uM LCA
10 uM P3865
10 uM Q1570
FF 1:300
MF 1:300
Ringers
ΔF/F
Frames after the beginning of each stimulus (3 sec/fr)
H
1 uM CA
1 uM CDCA
1 uM DCA
1 uM LCA
10 uM A6940
10 uM CA
10 uM CDCA
10 uM DCA
10 uM E0893
10 uM LCA
10 uM P3865
10 uM Q1570
FF 1:300
MF 1:300
Ringers
ΔF/F
Frames after the beginning of each stimulus (3 sec/fr)

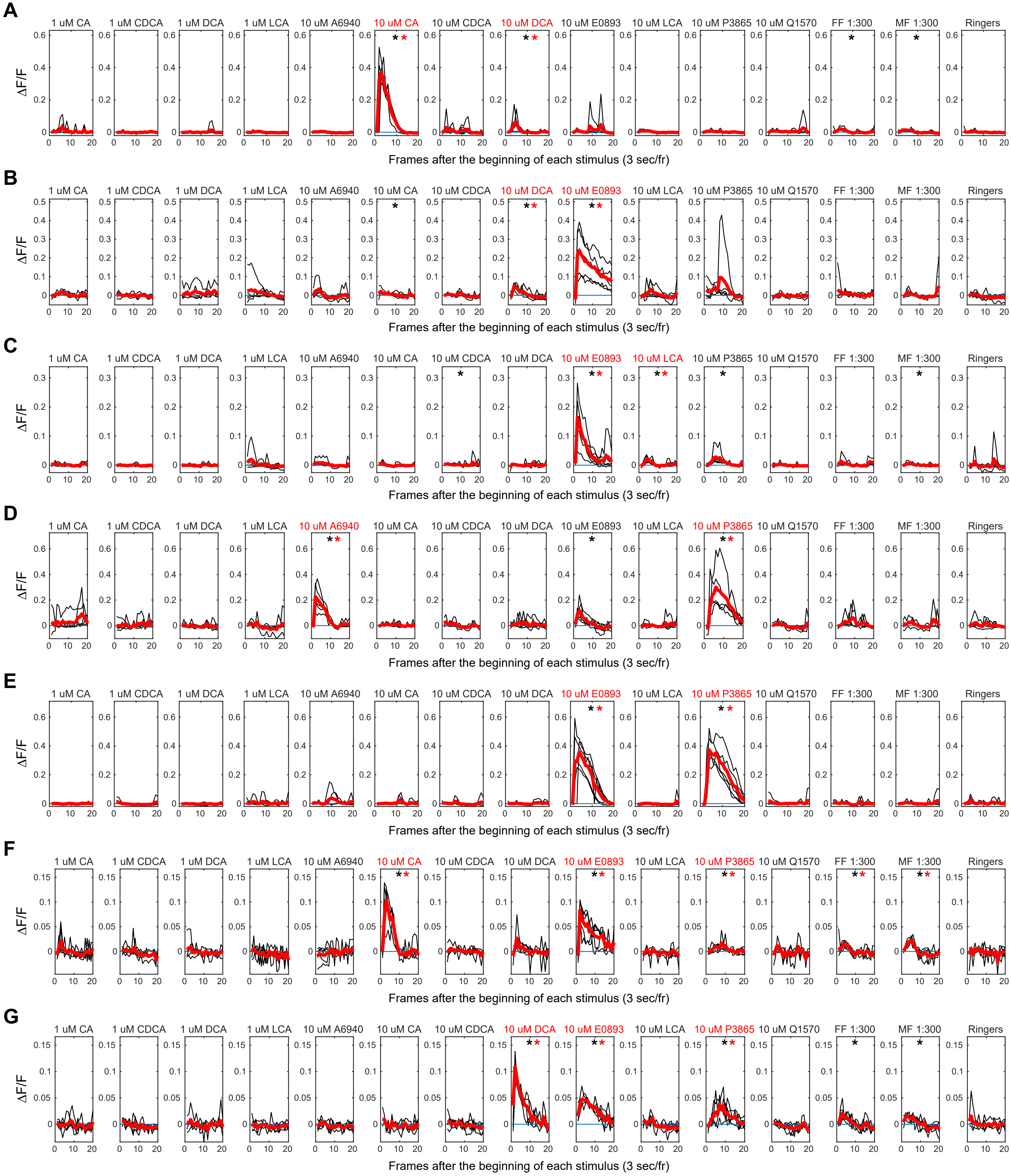

A
1 uM CA
1 uM CDCA
1 uM DCA
1 uM LCA
10 uM A6940
10 uM CA
10 uM CDCA
10 uM DCA
10 uM E0893
10 uM LCA
10 uM P3865
10 uM Q1570
FF 1:300
MF 1:300
Ringers
ΔF/F
Frames after the beginning of each stimulus (3 sec/fr)
B
1 uM CA
1 uM CDCA
1 uM DCA
1 uM LCA
10 uM A6940
10 uM CA
10 uM CDCA
10 uM DCA
10 uM E0893
10 uM LCA
10 uM P3865
10 uM Q1570
FF 1:300
MF 1:300
Ringers
ΔF/F
Frames after the beginning of each stimulus (3 sec/fr)
C
1 uM CA
1 uM CDCA
1 uM DCA
1 uM LCA
10 uM A6940
10 uM CA
10 uM CDCA
10 uM DCA
10 uM E0893
10 uM LCA
10 uM P3865
10 uM Q1570
FF 1:300
MF 1:300
Ringers
ΔF/F
Frames after the beginning of each stimulus (3 sec/fr)
D
1 uM CA
1 uM CDCA
1 uM DCA
1 uM LCA
10 uM A6940
10 uM CA
10 uM CDCA
10 uM DCA
10 uM E0893
10 uM LCA
10 uM P3865
10 uM Q1570
FF 1:300
MF 1:300
Ringers
ΔF/F
Frames after the beginning of each stimulus (3 sec/fr)
E
1 uM CA
1 uM CDCA
1 uM DCA
1 uM LCA
10 uM A6940
10 uM CA
10 uM CDCA
10 uM DCA
10 uM E0893
10 uM LCA
10 uM P3865
10 uM Q1570
FF 1:300
MF 1:300
Ringers
ΔF/F
Frames after the beginning of each stimulus (3 sec/fr)
F
1 uM CA
1 uM CDCA
1 uM DCA
1 uM LCA
10 uM A6940
10 uM CA
10 uM CDCA
10 uM DCA
10 uM E0893
10 uM LCA
10 uM P3865
10 uM Q1570
FF 1:300
MF 1:300
Ringers
ΔF/F
Frames after the beginning of each stimulus (3 sec/fr)
G
1 uM CA
1 uM CDCA
1 uM DCA
1 uM LCA
10 uM A6940
10 uM CA
10 uM CDCA
10 uM DCA
10 uM E0893
10 uM LCA
10 uM P3865
10 uM Q1570
FF 1:300
MF 1:300
Ringers
ΔF/F
Frames after the beginning of each stimulus (3 sec/fr)

**A**

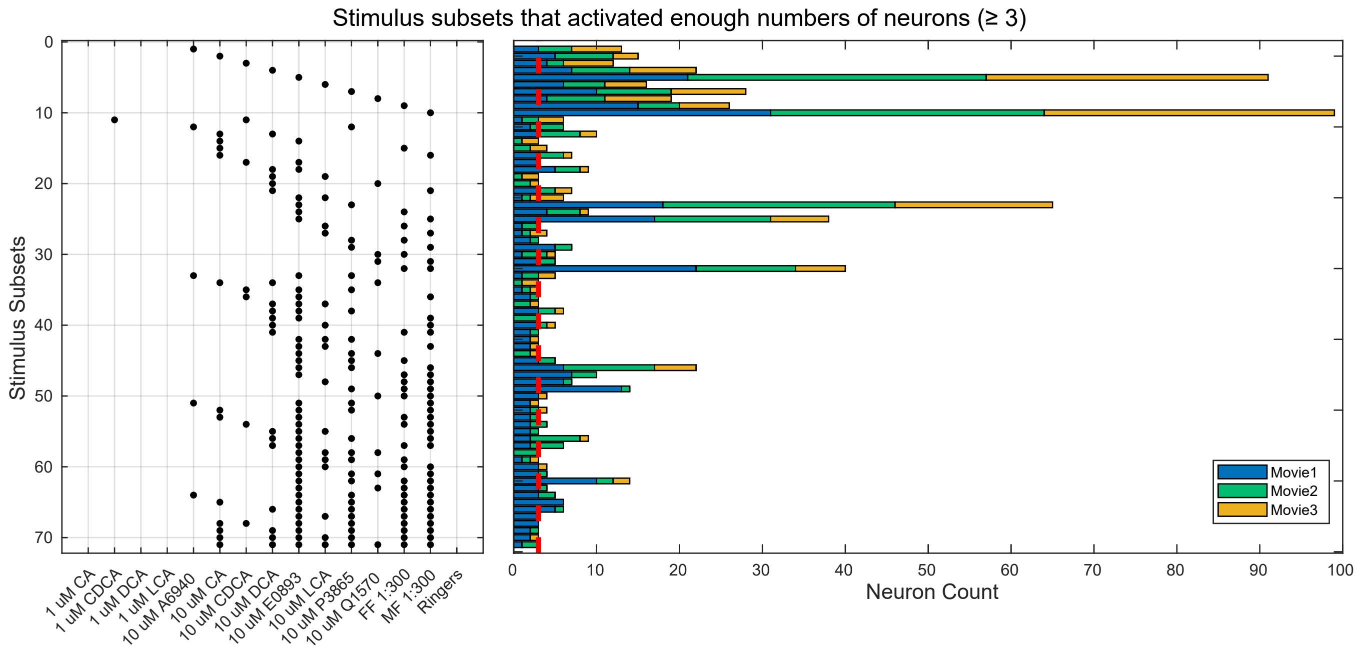


**B** Ligand subsets obtained from a threshold of 3

| | CA | DCA | LCA | CDCA | A6940 | E0893 | P3865 | Q1570 |
|---|---|---|---|---|---|---|---|---|
| 1 | ● | | | | | | | |
| 2 | | ● | | | | | | |
| 3 | | | ● | | | | | |
| 4 | | | | ● | | | | |
| 5 | | | | | ● | | | |
| 6 | | | | | | ● | | |
| 7 | | | | | | | ● | |
| 8 | | | | | | | | ● |
| 9 | ● | ● | | | | | | |
| 10 | ● | | | | | ● | | |
| 11 | | ● | ● | | | | | |
| 12 | | ● | | | | ● | | |
| 13 | | ● | | | | | | ● |
| 14 | | | ● | | | ● | | |
| 15 | | | | ● | | ● | | |
| 16 | | | | | ● | | ● | |
| 17 | | | | | | ● | ● | |
| 18 | | | | | | ● | | ● |
| 19 | ● | ● | | | | | | ● |
| 20 | ● | | | | | ● | ● | |
| 21 | | ● | ● | | | ● | | |
| 22 | | ● | | | | ● | ● | |
| 23 | | | ● | | | ● | ● | |
| 24 | | | | ● | | ● | ● | |
| 25 | | | | | ● | ● | ● | |
| 26 | | | | | | ● | ● | ● |
| 27 | ● | ● | | | | ● | ● | |
| 28 | ● | | | ● | | ● | ● | |
| 29 | | | ● | | | ● | ● | ● |
| 30 | ● | ● | ● | | | ● | ● | |
| 31 | ● | ● | ● | | | ● | ● | ● |

**C** Ligand subsets obtained from a threshold of 5

| | CA | DCA | LCA | CDCA | A6940 | E0893 | P3865 | Q1570 |
|---|---|---|---|---|---|---|---|---|
| 1 | ● | | | | | | | |
| 2 | | ● | | | | | | |
| 3 | | | ● | | | | | |
| 4 | | | | ● | | | | |
| 5 | | | | | ● | | | |
| 6 | | | | | | ● | | |
| 7 | | | | | | | ● | |
| 8 | | | | | | | | ● |
| 9 | ● | ● | | | | | | |
| 10 | | ● | | | | ● | | |
| 11 | | ● | ● | | | | | |
| 12 | | | ● | | | ● | | |
| 13 | | | | | ● | | ● | |
| 14 | | | | | | ● | ● | |
| 15 | ● | | | | | ● | ● | |
| 16 | | ● | | | | ● | ● | |
| 17 | | | | | ● | ● | ● | |

**D** Ligand subsets obtained from a threshold of 9

| | CA | DCA | LCA | CDCA | A6940 | E0893 | P3865 | Q1570 |
|---|---|---|---|---|---|---|---|---|
| 1 | ● | | | | | | | |
| 2 | | ● | | | | | | |
| 3 | | | ● | | | | | |
| 4 | | | | ● | | | | |
| 5 | | | | | ● | | | |
| 6 | | | | | | ● | | |
| 7 | | | | | | | ● | |
| 8 | | | | | | | | ● |
| 9 | ● | ● | | | | | | |
| 10 | | ● | | | | ● | | |
| 11 | | | | | | ● | ● | |
| 12 | | ● | | | | ● | ● | |
